# CRISPR-based kinome-screening revealed MINK1 as a druggable player to rewire 5FU-resistance in OSCC through AKT/MDM2/p53 axis

**DOI:** 10.1101/2021.08.19.456939

**Authors:** Sibasish Mohanty, Pallavi Mohapatra, Omprakash Shriwas, Shamima Azma Ansari, Manashi Priyadarshini, Swatismita Priyadarsini, Rachna Rath, Mahesh Sultania, Saroj Kumar Das Majumdar, Rajeeb Kumar Swain, Rupesh Dash

**Author notes:** Authors contributed equally (Sibasish Mohanty, Pallavi Mohapatra). Corresponding authors, Rupesh Dash, Institute of Life Sciences, Nalco Square, Chandrasekharpur, Bhubaneswar-751023, Odisha, India.

## Abstract

Cisplatin, 5FU and docetaxel (TPF) are the most common chemotherapy regimen used for advanced OSCC. However, many cancer patients experience relapse, continued tumor growth, and spread due to drug resistance, which leads to treatment failure and metastatic disease. Here, using a CRISPR/Cas9 based kinome knockout screening, Misshapen-like kinase 1 (MINK1) is identified as an important mediator of 5FU resistance in OSCC. Analysis of clinical samples demonstrated significantly higher MINK1 expression in the tumor tissues of chemotherapy non-responder as compared to chemotherapy responders. The nude mice and zebrafish xenograft experiments indicate that knocking out MINK1 restores 5FU mediated cell death in chemoresistant OSCC. An antibody based phosphorylation array screen revealed MINK1 as a negative regulator of p53. Mechanistically, MINK1 modulates AKT phosphorylation at Ser473, which enables p-MDM2 (Ser 166) mediated degradation of p53. We also identified lestaurtinib as a potent inhibitor of MINK1 kinase activity. The patient derived TPF resistant cell based xenograft data suggest that lestaurtinib restores 5FU sensitivity and facilitates a significant reduction of tumor burden. Overall, our study suggests that MINK1 is a major driver of 5FU resistance in OSCC. The novel combination of MINK1 inhibitor lestaurtinib and 5FU needs further clinical investigation in advanced OSCC.

## Introduction

Majority of head and neck cancer is originated from mucosal epithelium collectively termed as Oral squamous cell carcinomas (OSCC) ^1^. It is the most prevalent neoplasm in developing country like India with approximately 80000 new cases diagnosed every year ^2^. Unfortunately, most of the patients present with advanced OSCC are without having any preclinical history of pre malignant lesions. The treatment modalities for advanced OSCC includes surgical removal of tumor followed by concomitant chemoradiotherapy. Neoadjuvant chemotherapy is frequently prescribed for surgically unresectable OSCC tumors ^3^. However, despite of having all these treatment modalities the 5-year survival rate of advanced tongue OSCC is less than 50%. Chemoresistance is one of the major causes of treatment failure in OSCC^4^. The chemotherapeutic regimen used for OSCC are cisplatin, 5FU and Docetaxel (TPF)^3^. Though chemotherapy drugs show initial positive response, tumor acquires resistance gradually and patients experience continued tumor growth and metastatic disease.

Reprogramming resistant cells to undergo drug induced cell death is a viable way to overcome drug resistance. This can be achieved by identifying the causative factors for acquired chemoresistance and discovering novel agents to target critical causative factors, which will restore drug-induced cell death in chemoresistant OSCC. Kinases, which transfer a reversible phosphate group to proteins, play important role in regulating several phenotypes of carcinogenesis including growth, proliferation, angiogenesis, metastasis and evasion of antitumor immune responses ^5^. There are approximately 538 known kinases in human which are known to regulate different kinase signaling. A few of them are also known to regulate drug resistance in HNSCC. A kinome study revealed, microtubule-associated serine/threonine kinase 1 (MAST1) is a major driver of cisplatin resistance in HNSCC. MAST1 inhibitor lestaurtinib efficiently sensitized chemoresistant cells to cisplatin. Overall, the study suggests that MAST1 is a viable target to overcome cisplatin resistance ^6^. Ectopic overexpression of receptor tyrosine kinase in HNSCC mediates acquired resistance against cetuximab. For example, hyper activation of AXL was observed in clinical samples those are resistant to cetuximab ^7^. It was also found that RAS-MAPK are key mediator for cetuximab resistance in OSCC ^8^. However, very limited studies are available about the kinases those mediate 5FU resistance in OSCC.

MINK1 belongs to germinal center kinase (GCK) family and it is involved in regulation of several important signaling cascades ^9^. Recently, it is reported that MINK1 can regulate the planner cell polarity, which is essential for spreading of cancer cells. The PRICKLE1 encodes the planner cell polarity protein that binds to MINK1 and RICTOR (a member in mTOR2 complex) and this complex regulates the AKT meditated cell migration. Selectively targeting either of MINK1, PRICKLE1 or RICTOR can significantly decrease the migration of cancer cell in breast carcinomas ^10^. Ste20-related kinase, misshapen (msn), a Drosophila homolog of MINK1 regulates embryonic dorsal closure through activation of c-jun amino-terminal kinase (JNK) ^11^.

The goal of this study is to find out the potential kinase(s) those are major driver(s) of 5FU resistance in OSCC, for which a CRISPR based kinome screening was employed on 5FU resistant OSCC lines. The top ranked protein MINK1 was selected for validation in multiple cell lines and patient derived cells. In addition to this, lestaurtinib was identified to inhibit MINK1 kinase activity, which can reverse 5FU mediated cell death in chemoresistant OSCC lines. Ultimately, we demonstrated that MINK1 regulates the p53 in 5FU resistant OSCC. MINK1 activates AKT by phosphorylation at Ser473, which phosphorylates MDM2 at Ser166, the later in turn triggers degradation of p53.

## Materials and Methods

### Cell culture

The human tongue OSCC lines (H357, SCC4 and SCC9) were obtained from Sigma Aldrich, sourced from European collection of authenticated cell culture. All OSCC cell lines were cultured and maintained in DMEM F12 supplemented with 10% FBS (Thermo Fisher Scientific), penicillin–streptomycin (Pan Biotech) and 0.5 ug/ml sodium hydrocortisone succinate. HEK 293T cells were maintained in DMEM supplemented with 10% FBS and penicillin–streptomycin (Pan Biotech).

### High Content Screening

1000 cells/ well were seeded in black flat bottom 96 well plate (Thermo Scientific™ Nunc) and divided into two experimental groups, one without 5FU treatment and the other with 5FU treatment. A CRISPR based kinome-wide screening was performed using a lentiviral sgRNA library (LentiArray™ Human Kinase CRISPR Library, Thermo Fisher Scientific, M3775) that knocks out 840 kinase and kinase related genes individually with total number of 3214 sgRNA constructs. Transduction of lentiviruses (MOI:2) containing pooled sg RNAs (up to 4) targeting each of 840 genes along with positive and negative control lentiviruses into individual wells was carried out in presence of polybrene (8μg/ml). At 48 hours post transduction, selection with puromycin (0.5μg/ml) was performed for next 2-3 days, followed by treatment of vehicle control and 5FU at sub lethal dose respectively in both groups for 48h. Finally, cells were stained with LIVE/DEAD™ Viability/Cytotoxicity Kit (Thermo Fisher Scientific Cat # L3224) and high content screening was performed using CellInsight CX7 High-Content Screening (HCS) Platform. The green fluorescence indicates the living cells and red fluorescence indicates dead cells. Images from 20 fields per well were acquired using 10X objective lens. Two different fluorescent channels (excitation wavelengths - 488nm and 561nm) were used for acquiring images. Image analysis was performed using the HCS Studio software. A threshold value for each channel was set once and used for the entire screening. To identify the cells, segmentation was done. Some of the clumped and poorly segmented cells were excluded from further analysis on the basis of area, shape and intensity. On the basis of intensity, number of live and dead cells were counted and an objective mask (blue lines in the images) was created around each cell. For positive control, Cas9 over expressing cells were transduced with lentiviruses expressing sgRNA targeting human hypoxanthine phosphoribosyltransferase 1 (HPRT1) (LentiArray™ CRISPR Positive Control Lentivirus, human HPRT, Thermo Fisher Scientific Cat # A32829). HPRT1 knockout cells shows resistance to 6-thioguanine (6TG) induced cell death. For negative control, Cas9 over expressing cells were transduced with lentiviruses expressing gRNA with no sequence homology to any region of the human genome (LentiArray™ CRISPR Negative Control Lentivirus, Thermo Fisher Scientific, Cat # A32327).

### Patient Derived Xenograft

BALB/C-nude mice (6-8 weeks, male, NCr-Foxn1nu athymic) were purchased from Hylasco Bio-Technology Pvt. Ltd. For xenograft model early passage of patient-derived cells (PDC2) established from chemo non-responder patient (treated with TPF without having any response) was considered. Two million cells were suspended in phosphate-buffered solution-Matrigel (1:1, 100 μl) and transplanted into upper flank of mice. The PDC MINK1WT cells were injected in right upper flank and PDC MINK1KO cells were injected in the left upper flank of same mice. These mice were randomly divided into 2 groups (n=5) once the tumors reached a volume of 50 mm^3^ and injected with vehicle control or 5FU (10mg/kg) intraperitonially twice a week. In another experimental set up, PDC2 WT cells were injected in right upper flank of mice. These mice were randomly divided into 4 groups (n=5) after the tumors have reached a volume of 50 mm^3^ and injected with vehicle control, 5FU (10mg/kg), Lestaurtinib (20mg/kg) and 5FU (10mg/kg) and Lestaurtinib (20mg/kg) respectively in each individual group, intraperitonially twice a week. Tumor size was measured using digital Vernier caliper twice a week until the completion of experiments. Tumor volume was determined using the following formula: Tumor volume (mm^3^) = (minimum diameter)^2^ × (maximum diameter)/2.

### Phospho-Protein Profiling

For performing the Phospho Explorer Antibody Array (Full Moon Biosystems Cat# PEX10), 5X10^6^ number of H3575FUR MINK1WT and H3575FUR MINK1KO cells were seeded and lysates were isolated, after which the protein samples were labelled with biotin as described by manufacturer. Biotinylated protein samples were further blocked in skim milk and were subjected to coupling with the 1318 number of antibodies present on the array slides. Then for the detection of expression of phospho proteins, Cy3-streptavidin (Sigma, Cat#S6402) was added. Array slides were scanned at Fullmoon Biosystems array scanning service and the image analysis was done using ImageJ software.

### *In vitro* MINK1 kinase activity assays

MINK1 Kinase Enzyme System (Promega Cat No# V3911) and ADP-Glo™ Kinase Assay (Promega Cat No#V9101) were procured to perform the *in vitro* kinase assay. Selected compounds (10 μM) were incubated at room temperature for 1 hr with recombinant human MINK1 kinase along with substrate MBP and ATP to perform the kinase reaction using 1X kinase buffer. Further ADP-Glo™ Reagent was added at room temperature for 40 mins to stop the kinase reaction and deplete the unconsumed ATP, hence leaving only ADP. Then Kinase detection reagent was added and incubated at room temperature for 30 mins to convert ADP to ATP and introduce luciferase and luciferin to detect ATP. Finally, the luminescence was measured using VICTOR® Nivo™ Multimode Plate Reader (Perkin Elmer).

### Statistical analysis

All data points are presented as mean and standard deviation and Graph Pad Prism 9.0 was used for calculation. The statistical significance was calculated by one-way variance (one-way ANOVA), Two-Way ANOVA and considered significance at P≤0.05.

### Study approval

This study was approved by the Institute review Board and Human Ethics committees (HEC) of Institute of Life Sciences, Bhubaneswar (110/HEC/21) and All India Institute of Medical Sciences (AIIMS), Bhubaneswar (T/EMF/Surg.Onco/19/03). The animal related experiments were performed in accordance to the protocol approved by Institutional Animal Ethics Committee of Institute of Life Sciences, Bhubaneswar (ILS/IAEC-204-AH/DEC-20). Approved procedures were followed for patient recruitment and after receiving written informed consent from each patient, tissues samples were collected. Institutional biosafety committee (IBSC) approved all related experiments.

## Results

### Establishment and characterization of 5FU resistant OSCC lines

The 5FU resistant OSCC lines were established by prolonged treatment of 5FU to OSCC cell lines as described in materials and methods. Monitoring the cell viability of 5FU sensitive (5FUS) and resistant (5FUR) pattern of H357, SCC4 and SCC9 cell lines by MTT assay suggest that 5FUR cells achieved acquired resistance (Fig. S1A,B). Enhanced cancer like stem cells (CSCs) and elevated expression of ATP-binding cassette (ABC) transporters are the hallmarks of chemoresistant cells. qRT-PCR data suggest that CSC markers (SOX2, OCT4 and NANOG) as well as majority of ABC transporters expression were elevated in 5FUR cells as compared to 5FUS cells (Fig. S1C, D).

### Kinome wide screening identifies MINK1 as a potential driver of 5FU resistance in OSCC

To identify the kinases those play important role in 5FU resistance, a CRISPR based kinome-wide screening was performed using a lentiviral sgRNA library knocking out 840 kinases individually with a total number of 3214 sgRNA constructs. To target the individual kinase, upto 4 sgRNA lentiviral constructs were pooled together. For kinome screening, Cas9 overexpressing 5FU resistant OSCC lines were established (Fig. S2), which showed similar drug resistant pattern with parental 5FUR OSCC lines(Fig. S3A). We also determined the polybrene and puromycin tolerance concentration in Cas9 overexpressing clones (Fig. S3B, C). The 5FUS and 5FUR lines were treated with 5FU and cell death was measured in high content analyzer using a fluorescent cell viability dye, the data suggest significantly lower cell death in 5FUR cells as compared to 5FUS cells, which is in harmony with the previously measured 5FU tolerance in both lines (Fig. S4A-C). The screening protocol was optimized using appropriate positive and negative control. When 6TG (6-Thioguanine) was treated to HPRT1 (Hypoxanthine Phosphoribosyltransferase 1) KO lines, which is used as positive control for screening, it showed resistance to cell death, whereas HPRT1 WT cells were sensitive to 6TG, suggesting optimized screening protocol (Fig. S4D, E). As a negative control, lentivirus expressing scrambled sgRNA was used.

For primary screening, the 5FU resistant line (H357 5FUR) was transduced with a lentivirus containing sgRNAs targeting each of the 840 individual kinases, after which sub lethal dose of 5FU was treated for 48h followed by measuring cell death in high content analyzer using a fluorescent cell viability dye (Fig. 1A). From primary screening, 334 kinases out of 840 were selected for further consideration by rejecting rest of sgRNA clones which alone induced cell death more than 30 % (Fig. 1B, C). The 60 candidate kinases having lowest survival fraction score were evaluated in the secondary screening using three more chemoresistant lines i.e SCC4 5FUR, SCC9 5FUR and H357 CisR. From the primary and secondary screening, MINK1, SBK1 and FKBP1A emerged as the only three common kinases among the 5FUR lines with MINK1 having the lowest survival fraction score, which sensitize the chemoresistant cell to 5FU mediated cell death the most (Fig. 1 C-E). In secondary screening, MINK1 knock out showed minimal efficacy in sensitizing cisplatin resistant cell lines to cisplatin (Fig. 1D), indicating the specific role of MINK1 towards acquired 5FU resistance. Next, monitoring the expression of MINK1 in OSCC resistant lines, we found the expression of MINK1 is significantly higher in 5FUR lines as compared to 5FUS lines of OSCC (Fig. 1F). With the evaluation of clinical samples, the expression of MINK1 was found to be elevated in tumor tissues of chemotherapy non-responders as compared to chemotherapy responders (Fig. G, H). We also evaluated the MINK1 expression in drug-naive and post-CT nonresponder paired tumor samples from the same patient and observed that the post–CT-treated tumor samples showed higher MINK1 expression (Fig. I, J).

**Figure 1:**
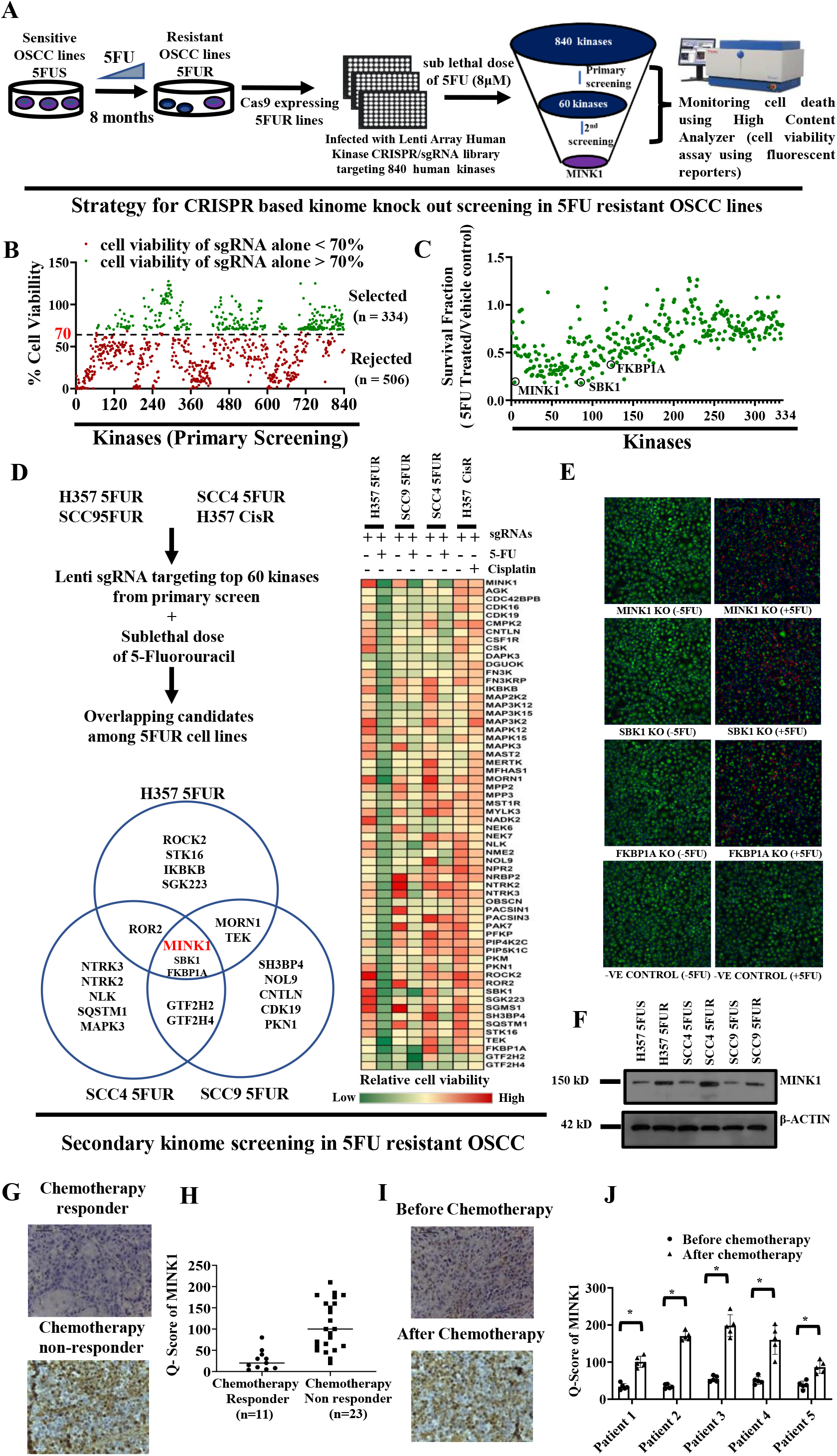
CRISPR based Kinome screening revealed MINK1 as a potential mediator for 5FU resistance in OSCC. **A)** Schematic presentation of approach for CRSPR/Cas9 based kinome knockout screening to discover the potential kinase responsible for 5FU resistance in OSCC. **B-C)** Primary screening of 840 kinases was performed with sublethal dose of 5FU (8μM). The kinases (n=506 nos) whose knockout alone induced significantly higher cell death (> 30%) depicted in red were excluded. From the rest of the kinases (n=334 nos) depicted in green, the survival fraction (5FU treated/ Vehicle Control) was determined and top 60 candidates having lowest survival fraction were considered for secondary screening. **D)** For secondary screening with top 60 kinases, four cell lines were considered i.e., H357 5FUR, SCC4 5FUR, SCC9 5FUR and H357CisR. After overlapping all three 5FUR cell lines, MINK1, SBK1, FKBP1A were found to be the common kinases among them. MINK1 was selected as a potential kinase target purely based on having the lowest survival fraction among all common candidates. **E)** The fluorescent images acquired from high content analyzer with indicated treated group during kinome screening. **F)** Lysates were collected from indicated cells and immunoblotting was performed with indicated antibodies. **G)** Protein expression of MINK1 was analyzed by IHC in chemotherapy-responder and chemotherapy-non-responder OSCC tumors. Scale bars: 50 μm. **H)** IHC scoring for MINK1 from panel G (Q Score =Staining Intensity × % of Staining), (Median, n=11 for chemotherapy-responder and n=23 for chemotherapy-non-responder) \**P* < 0.05 by 2-tailed Student’s t test. **I)** Protein expression of MINK1 was analyzed by immunohistochemistry (IHC) in pre- and post-TPF treated paired tumor samples from chemotherapy-non-responder patients. Scale bars: 50 μm. **J)** IHC scoring for MINK1 from panel I (Q Score =Staining Intensity × % of IHC Staining). \**P* < 0.05 by 2-tailed Student’s t test.

### MINK1 is an important target to overcome 5FU resistance in OSCC

To confirm the finding from the kinome screening, MINK1KO (knock out) clones were generated, using lentivirus expressing two different sgRNAs, in Cas9 overexpressing 5FUR OSCC lines and patient derived line 2 (PDC2) (Fig. S5A). PDC2 was isolated and characterized earlier from tumor of chemotherapy-non-responder patient, who was treated with neoadjuvant TPF without any response ^22^. The colony forming, MTT and spheroid assay data suggest that knocking out MINK1 significantly reduced the cell viability when the chemoresistant cells were treated with 5FU (Fig. S5B, C and Fig.2A). Here onwards we used sgRNA 1 for rest of the experiments. Similarly, knocking out MINK1 induced 5FU mediated cell death in chemoresistant cells (Fig. 2B). Enhanced p-H2AX and cleaved PARP was observed in MINK1KO cells followed by treatment with 5FU indicating the potential role of MINK1 in mediating 5FU resistance (Fig. 2C,D). Further, to test the in vivo efficacy of the kinome screening data, we implanted PDC2 MINK1WT cells into right upper flank and PDC2 MINK1KO cells into the left upper flank of nude mice followed by treatment with 5FU. Treatment with 5FU (10 mg/kg) significantly reduced the tumor burden in the MINK1KO but not in MINK1WT group (Fig. 2E-G). Immunohistochemistry data suggests markedly decreased cell proliferation signal (Ki67) in 5FU-treated MINK1KO tumors (Fig. 2H). Earlier it is known that selective knockdown of MINK1 decreases the migration of human breast cancer lines ^10^. To evaluate whether depletion of MINK1 also reduces migration of chemoresistant OSCC lines and PDC2, Boyden chamber assays and scratch/wound healing assays were performed. The data suggest that knock out of MINK1 followed by treatment with 5FU significantly reduces the relative number of migrated cells (Fig. S6A). Similarly, scratch area analysis suggest that percentage of scratch area is significantly higher when 5FU is treated to MINK1KO drug resistant cells (Fig. S6B, C). Next, we used Zebrafish (Danio rerio) [Tg(fli1:EGFP)] tumor xenograft model to further validate our findings. Equal number of the WT and PDC2MINK1 KO cells were stained with Dil (1,1’-Dioctadecyl-3,3,3’,3’-Tetramethylindocarbocyanine Perchlorate) and injected into perivitelline space of 48-hour post fertilized zebrafish embryos. After 3 days of injection, embryos were treated with vehicle control or 5FU (500μM). After 5 days of injection, the primary tumors and metastatic distribution of cancer cells were documented using a fluorescence microscope. The tumor growth, as measured by fluorescence intensity of primary tumors, was found to be significantly reduced in the MINK1 KO group with the treatment of 5FU (Fig. 2I, J). Also, cancer cells showed reduced distal migration from the primary site in the case of MINK1 KO 5FU treated group (Fig. 2K). These data indicate MINK1 dependency of 5FU resistant OSCC.

**Figure 2:**
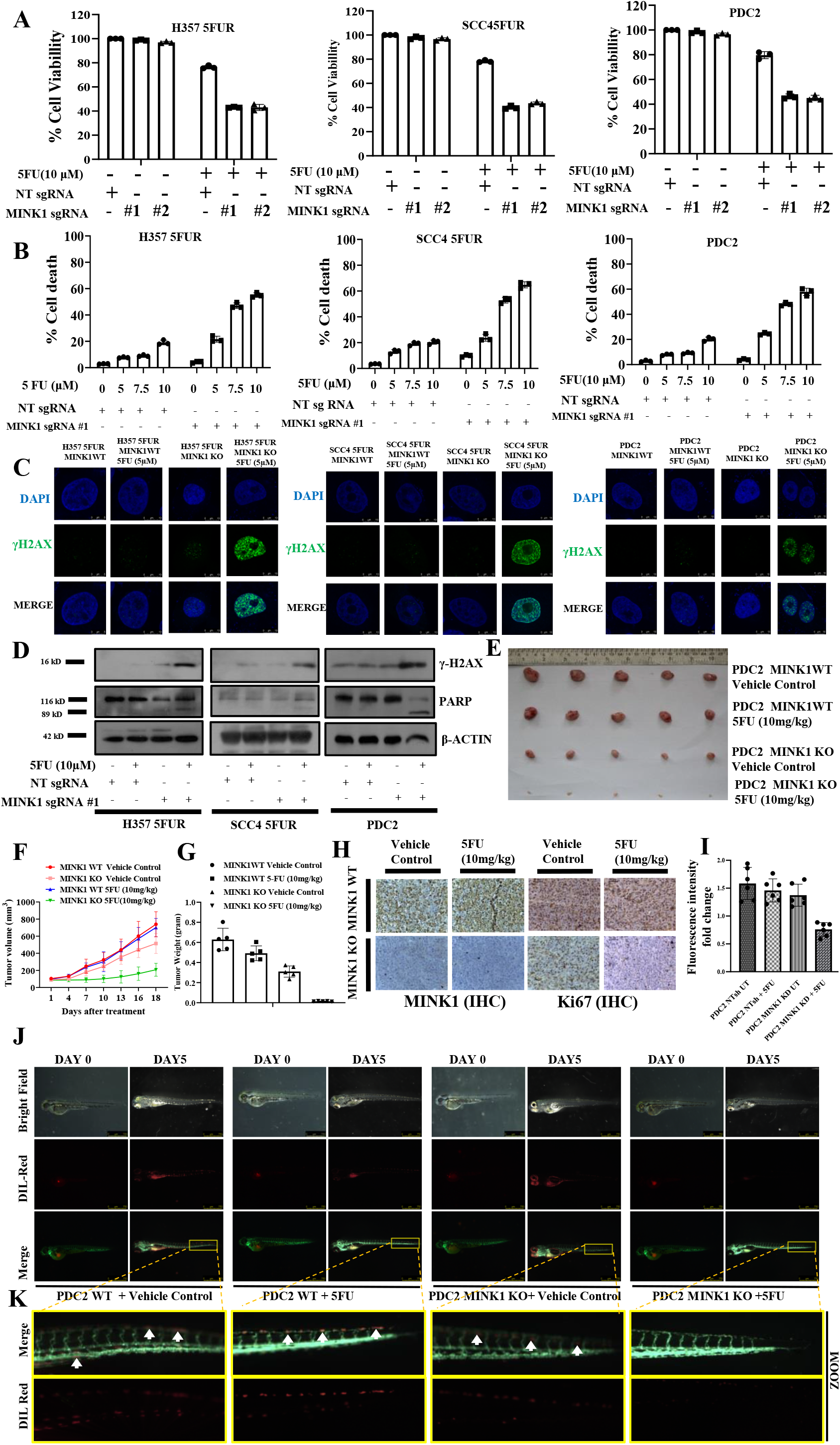
Selectively targeting MINK1 restores 5FU induced cell death in chemoresistant OSCC: **A)** 5FU resistant cells stably expressing MINK1sgRNA (#1 and #2) and NTsgRNA were treated with 5FU for 48h and cell viability was determined by MTT assay (n=3 and 2-way ANOVA). **B)** Indicated cells were treated with 5FU for 48h, after which cell death was determined by annexin V/7AAD assay using flow cytometer. Bar diagrams indicate the percentage of cell death (early and late apoptotic) with respective treated groups (Mean ±SEM, n=3, Two-way ANOVA). **C)** Indicated cells were treated with 5 μM of 5FU for 48h, after which immunostaining was performed for γ-H2AX. **D)** Indicated cells were treated with 5FU for 48h and immunoblotting was performed with indicated antibodies. **E)** PDC2 MINK1WT cells were implanted in right upper flank of athymic male nude mice and PDC2 MINK1KO cells were implanted in left upper flank, after which they were treated with 5FU at indicated concentration. At the end of the experiment mice were euthanized, tumors were isolated and photographed (n=5). **F)** Tumor growth was measured in indicated time points using digital slide caliper and plotted as a graph (mean ± SEM, n = 5). Two-way ANOVA. **G)** Bar diagram indicates the tumor weight measured at the end of the experiment (mean ± SEM, n = 5). Two-way ANOVA. **H)** After completion of treatment, tumors were isolated and paraffin-embedded sections were prepared as described in materials and methods to perform immunohistochemistry with indicated antibodies. Scale bars: 50 μm. **I, J)** Lateral view of fluorescent transgenic [Tg(fli1:EGFP)] zebrafish embryos at Day 0 and Day 5 injected with Dil-Red stained PDC2 control and MINK1 KO cells with and without treatment of 5FU **(J)**. The tumor growth and migration was assessed by an increase in fluorescence intensity on the 5th day compared to the day of injection. n=6. The quantitation of fluorescence intensity **(I)** was performed using ImageJ software and represented as fold change of fluorescence intensity where day 0 reading was taken as baseline. **K)** Zoomed image of distal part of embryo (5days post injection) to monitor migration of tumor cells.

### Ectopic expression of MINK1 promotes 5FU resistance in OSCC

To confirm the potential role of MINK1 in 5FU resistance, we performed gain of function study. For this, using a lentiviral approach we generated MINK1ShRNA stable clones in 5FUR lines and PDC2 (MINK1UTRKD), where the shRNA targets the 3’UTR of MINK1 mRNA. For ectopic overexpression of MINK1, the MINK1UTRKD cells were transfected with pDESTCMV/TO MINK1 vector (Fig. 3A). The cell viability and cell death data suggest that knocking down MINK1 in 5FUR cells result in sensitizing the resistant cells to 5FU, however ectopic overexpression of MINK1 rescues the 5FU resistant phenotype (Fig. 3B, C). Similarly, immunostaining data suggest enhanced p-H2AX signal in MINK1UTRKD cells, whereas ectopic overexpression of MINK1 reduces the p-H2AX signal indicating rescue of 5FU resistance in OSCC cells (Fig. 3D). We also observed the rescue of cleaved PARP with ectopic expression of MINK1 suggesting reduced cell death (Fig. 3E). Finally, when MINK1 was overexpressed in OSCC sensitive lines, cells showed resistance to 5FU induced cell death (Fig. 3F).

**Figure 3:**
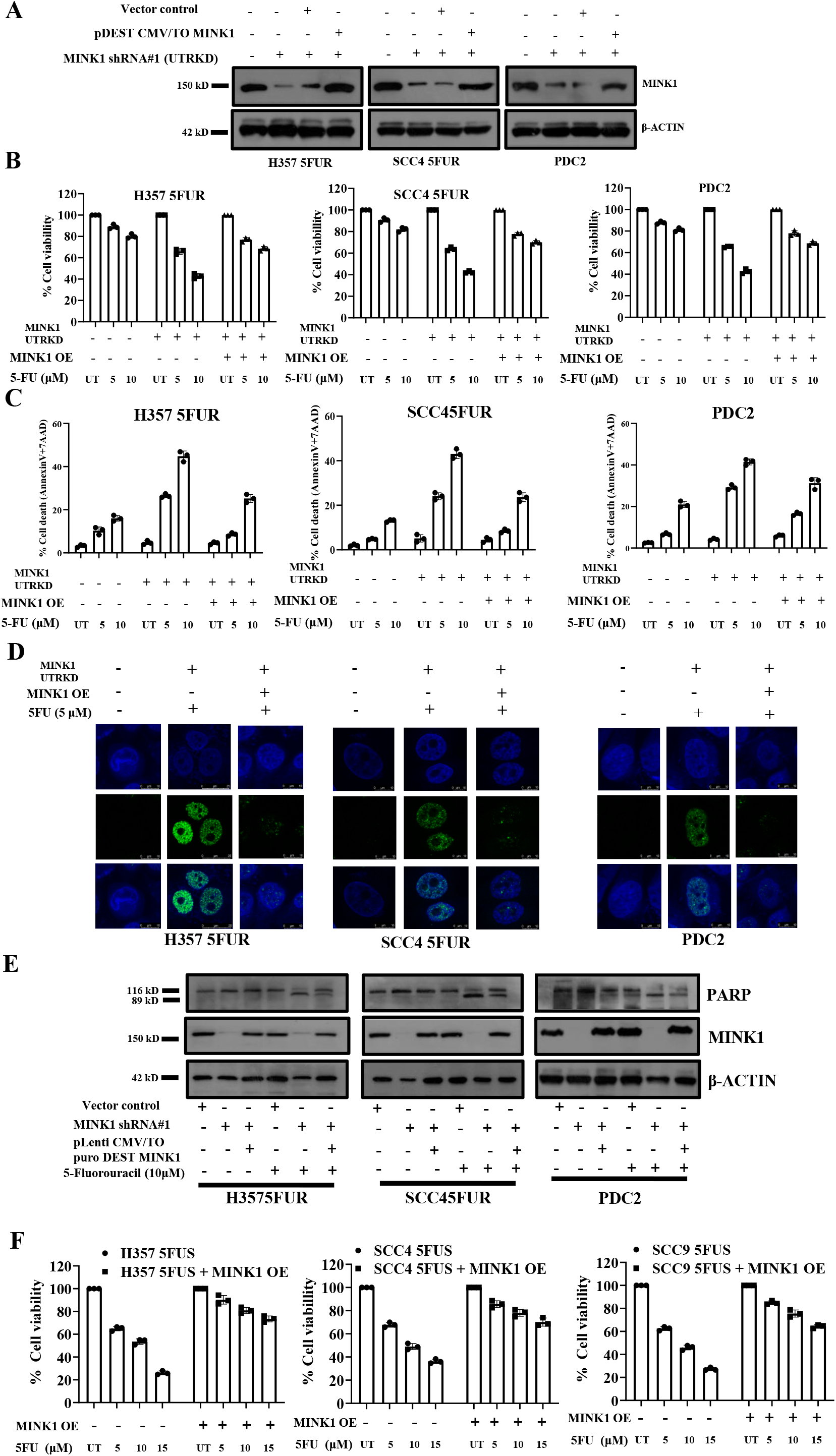
Ectopic overexpression of MINK1 rescued the drug resistant phenotype in MINK1KD drug resistant OSCC: **A)** Using a lentiviral approach, 5FU resistant OSCC lines and PDC2 were stably transfected with ShRNA which targets 3’UTR of MINK1 mRNA (MINK1 UTRKD). For ectopic overexpression, pLenti CMV/TO Puro DEST MINK1 and control vector were transiently transfected to indicated MINK1 UTRKD cells and immunoblotting (n=3) was performed with indicated antibodies. **B)** MINK1 was ectopically overexpressed in 5FUR cells stably expressing MINK1ShRNA targeting UTR and treated with 5FU at indicated concentration for 48 h, after which cell viability was determined by MTT assay (n=3), 2-way ANOVA. **C)** Cells were treated as indicated in B panel and cell death (early and late apoptotic) was determined by annexin V/7AAD assay using flow cytometer. Bar diagrams indicate the percentage of cell death with respective treated groups (Mean ±SEM, n=3), 2-way ANOVA. **D)** MINK1 was overexpressed in 5FUR cells stably expressing MINK1ShRNA targeting UTR and treated with 5FU for 48h, after which immunostaining was performed for γ-H2AX as described in materials and methods. **E)** MINK1 was overexpressed in chemoresistant cells stably expressing MINK1ShRNA targeting 3’ UTR, followed by 5FU treatment for 48 hours, and immunoblotting (n = 3) was performed with indicated antibodies. **F)** 5FU sensitive OSCC lines were transfected with pLenti CMV/TO Puro DEST MINK1 followed by treatment with 5FU at indicated concentration for 48h, after which cell viability was determined by MTT assay (n=3), 2-way ANOVA.

### MINK1 downregulates the expression of p53 in chemoresistant OSCC through activation of AKT and MDM2

To understand the specific role of MINK1 in 5FU resistant OSCC, we performed high-throughput phosphorylation profiling with 1,318 site-specific antibodies from over 30 signaling pathways in 5FUR cells stably expressing MINK1KO and MINK1WT. From this study, phosphorylation of p53 at Ser33 and Ser15 were found to be significantly up regulated in MINK1KO cells as compared to MINK1WT cells. In addition to this, phosphorylation of AKT at Ser473 and phosphorylation of MDM2 at Ser166 were found to be down regulated in MINK1KO cells as compared to MINK1WT (Fig. 4A). Further, immunoblotting was performed to validate the finding of phosphorylation profiling antibody array. The data suggest that phosphorylation of p53 at Ser15 and Ser33 is significantly upregulated and phosphorylation of MDM2 at Ser166 is profoundly downregulated in MINK1KO cells as compared to MINK1WT chemoresistant cells (Fig. 4B). OSCC lines H357 and SCC4 have mutant p53, whereas MCF7 and HEK 293 have wild type p53 expression. Further, p53 expression was also found to be inversely correlated with MINK1 irrespective of its mutation status. (Fig. 4B, S7). Next, when MINK1 was ectopically overexpressed in MINK1KD (shRNA targeting 3’UTR) clones, downregulation of p53, p-p53 (Ser33) and p-p53 (Ser15) were observed in chemoresistant OSCC lines (Fig. 4C). In harmony to our finding of phosphorylation array, p-AKT(Ser473) was found to be down regulated in MINK1 depleted cells, which was rescued with ectopic overexpression of MINK1 (Fig. 4 D, E). To confirm the potential role of AKT in modulating MINK1 mediated p53 regulation, we ectopically overexpressed constitutively active AKT (myrAKT) in MINK1KO cells. The immunoblotting data suggest that expression of p53, p-p53 (Ser33) and p-p53 (Ser15) were downregulated when MyrAKT was overexpressed in MINK1KO cells (Fig. 4F). Similarly, when MINK1 over expressing cells were treated with AKT inhibitor (Akti-1/2), the p53 expression was rescued along with downregulation of p-MDM2 (Ser166) (Fig. 4G). p53 target genes were also evaluated in MINK1 KO cells and the immunoblotting data suggest that expression of p21, NOXA and TIGAR in MINK1KO clones were upregulated as compared to MINK1WT clones (Fig. 4H).

**Figure 4:**
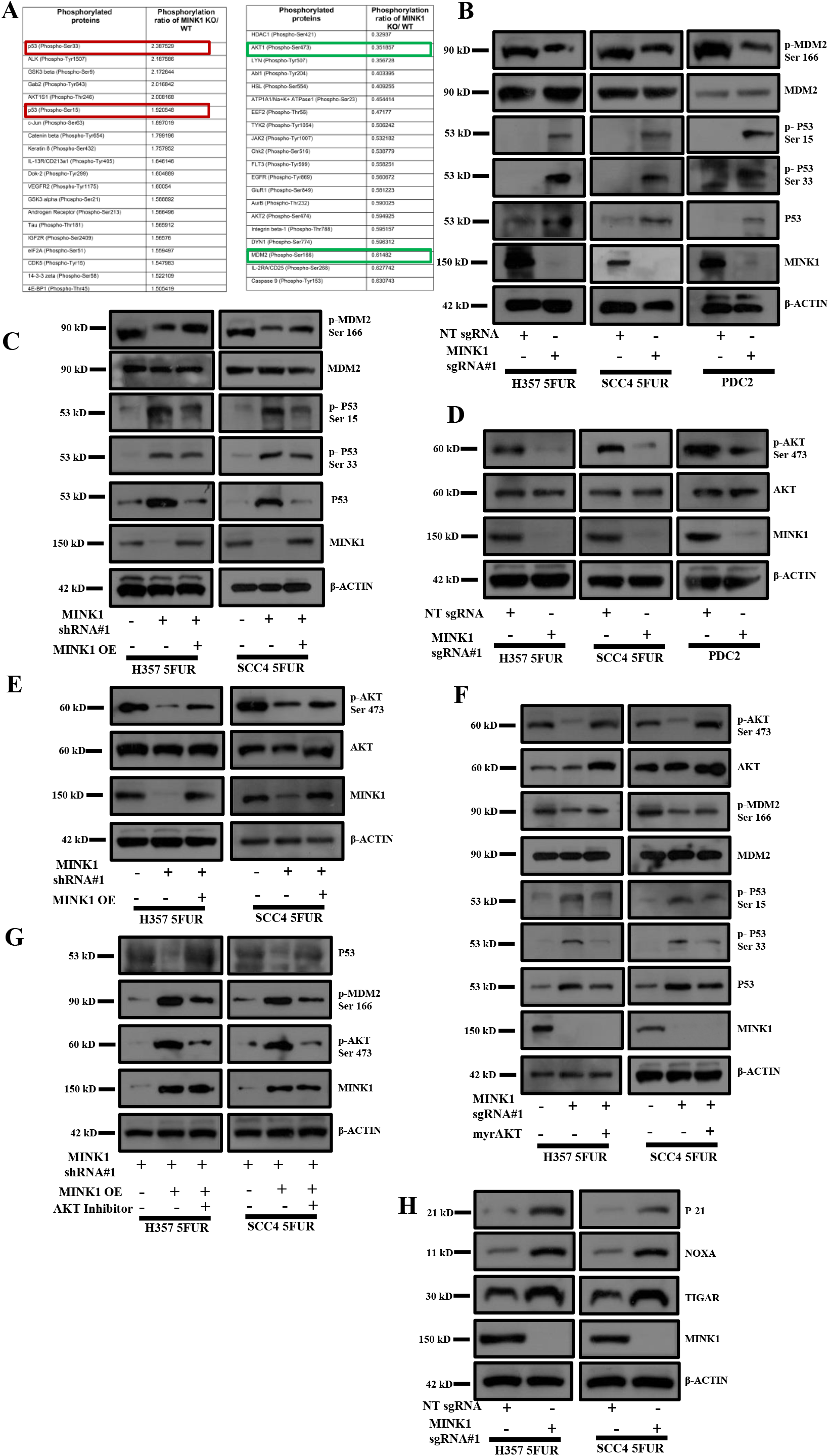
MINK1 regulates the expression of p53 in 5FU resistant OSCC lines through AKT/MDM2. **A)** High-throughput phosphorylation profiling with 1,318 site-specific antibodies from over 30 signaling pathways was performed in the lysates of MINK1KO and MINK1WT clones of H357 5FUR cells as described in methods. The top 20 upregulated phosphoproteins (MINK1 KO/MINK1WT) is represented in left panel, whereas top 20 downregulated phosphorylated proteins is represented in right panel. The upregulated targets considered in the study is marked in green red box, whereas downregulated targets in green box. **B)** Lysates were collected from indicated cells and immunoblotting was performed with indicated antibodies. **C)** pDESTCMV/TO MINK1 (ectopic overexpression of MINK1) was transiently transfected in 5FUR lines stably expressing MINK1 ShRNA (targeting 3’UTR) and immunoblotting was performed with indicated antibodies. **D)** Lysates were collected from indicated cells and immunoblotting was performed with indicated antibodies. **E)** MINK1 was ectopically overexpressed in 5FUR lines stably expressing MINK1 ShRNA (targeting 3’UTR) and immunoblotting was performed with indicated antibodies. **F)** pLNCX myr HA Akt1 (ectopic overexpression of myr AKT) was transiently transfected in indicated MINK1KO cells and immunoblotting was performed with indicated antibodies. **G)** MINK1 was ectopically overexpressed in 5FUR lines stably expressing MINK1ShRNA (UTRKD) as described in panel C and treated with AKT inhibitor (Akti-1/2) for 24h and immunoblotting was performed with indicated antibodies. **H)** Lysates were collected from indicated cells and immunoblotting was performed with indicated antibodies.

### Evaluation of Lestaurtinib as MINK1 inhibitor to reverse 5FU resistance in OSCC

From the screening data, we observed that MINK1 expression is elevated in chemoresistant OSCC and genetic inhibition of the same sensitizes drug resistant lines to 5FU induced cell death. Hence, MINK1 can be a potential therapeutic target to overcome chemoresistance in OSCC. Very limited information on the inhibitors of MINK1 is available in the literature. Hence, we looked for the potential MINK1 inhibitors in the international union of basic and clinical pharmacology (IUPHAR) database, where a screen of 72 inhibitors against 456 human kinases binding activity is provided. Among the potential twelve MINK1 inhibitors, we tested the MINK1 inhibitory activity of three inhibitors i.e., staurosporine, pexmetinib and lestaurtinib. The kinase assay data suggest that lestaurtinib and pexametinib have highest inhibitory activity for MINK1 (Fig. 5A). The 50% MINK1 inhibitory activity was observed at concertation of 100 nM in case of lestaurtinib and 10 μM for pexmetinib (Fig. 5B). Next, cell viability assay was performed to select a dose of lestaurtinib and pexmetinib that does not affect cell viability when treated alone (viability > 80%) in 5FU resistant OSCC lines (Fig. 5C, D). Further, the cell viability, spheroid assay and cell death data suggest that the selected sub lethal dose of lestaurtinib (50nM) and pexmetinib (500μM) can efficiently restore 5FU mediated cell death in chemoresistant OSCC lines and PDC2 (Fig. 5 E, F and S8A). The IC50 value of 5FU in H3575FUR is 20.49μM, however combination of lestaurtinib (50nM) decreases the IC50 value to 4.82 μM and combination of pexmetinib (500μM) lowers the IC50 value of 5FU to 7.08 μM (Fig. 5 E). As lestaurtinib, with a much lower concentration (50nM) as compared to pexmetinib (500nM) sensitize 5FU to chemoresistant cells, from here on lestaurtinib was considered for rest of the study. Enhanced expression of p-H2AX and cleaved PARP was observed only in combination group with lestaurtinib and 5FU indicating programmed cell death (Fig. 5G and S8B). Boyden chamber assays data suggests that combinatorial treatment of lestaurtinib and 5FU significantly reduces the relative number of migrated cells (Fig S8C). In harmony to the observation made by knockout of MINK1 in chemoresistant cells, we also found that lestaurtinib significantly decreased the phosphorylation of MDM2(Ser166) and AKT(Ser473) and elevated the expression of p53 in chemoresistant cells (Fig. 5H). Further, we found that lestaurtinib failed to sensitize 5FU mediated cell death in MINK1 knocked out 5FUR lines (Fig. 5I), which suggests that lestaurtinib conferred 5FU sensitivity by inhibiting MINK1 kinase activity. To check the in vivo efficacy of this novel combination, nude mice xenograft model was performed using patient derived cells (PDC2). The in vivo data suggest that the combination of lestaurtinib (20mg/kg) and 5FU (10mg/kg) profoundly reduced the tumor burden as compared to treatment with either of the single agents (Fig. 6A-C). Immunohistochemistry data suggest significant reduction in CD44 and Ki67 expression along with increased expression of cleaved caspase 3 in combination group (Fig. 6D). Finally, we performed combinatorial anti-tumor effect of non-cytotoxic extremely low dose of cisplatin (1μM), 5FU (1μM) and lestaurtinib (50nM) in PDC2. The cell viability, cell death, western blotting and colony forming assay data suggest significantly higher cell death in cisplatin, 5FU and lestaurtinib combinatorial group, as compared to any other possible combinatorial group, i.e. 5FU and lestaurtinib or cisplatin and lestaurtinib or cisplatin and 5FU (Fig. S8).

**Figure 5:**
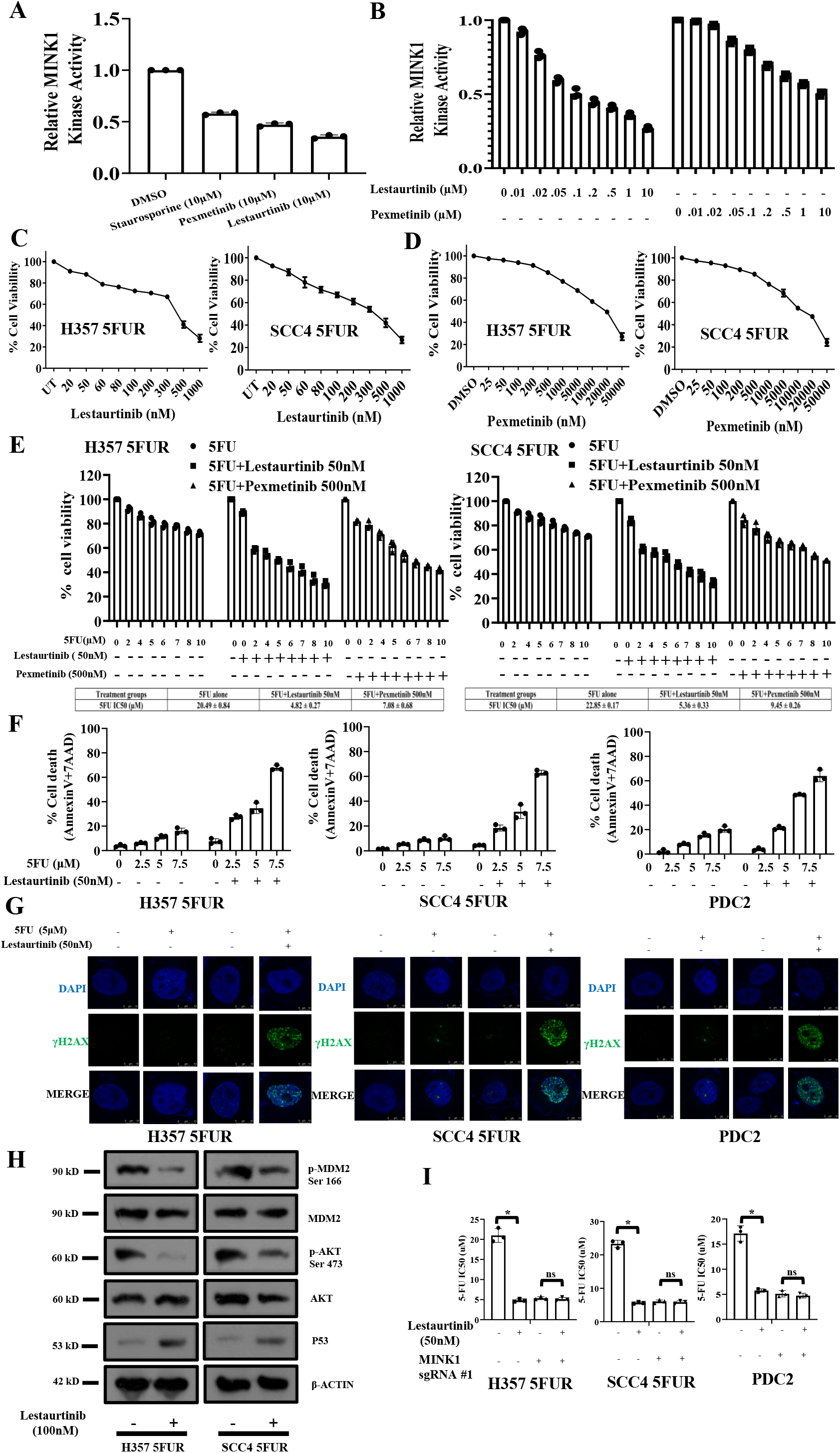
Evaluation of Lestaurtinib as a MINK1 inhibitor to restore 5FU sensitivity in drug resistant OSCC: **A)** *In vitro* MINK1 kinase assay was performed using three compounds potentially binding to MINK1 (based on IUPHAR database). All compounds (10 μM) were incubated with recombinant human MINK1 along with substrate MBP and ATP and further subjected to ADP-Glo™ Kinase Assay as described in materials and methods section. **B)** Determination of EC50 value for kinase activity of top two MINK1 inhibitors selected from panel (A). **C-D)** Selection of highest dose of Lestaurtinib and Pexmetinib that does not affect cell viability when treated alone (viability > 80%) in 5FU resistant OSCC lines (n=3), 2-way ANOVA. **E)** 5FU resistant cells were treated with indicated dose of MINK1 inhibitor (50 nM Lestaurtinib, 500 nM Pexmetinb) in combination with increasing concentrations of 5FU for 48 h, after which cell viability was determined by MTT assay (n=3), 2-way ANOVA. **F)** 5FU resistant OSCC lines and PDC2 cells were treated indicated dose of MINK1 inhibitor (50 nM Lestaurtinib, 500 nM Pexmetinb) in combination with increasing concentrations of 5FU for 48 h, after which cell death (early and late apoptotic) was determined by annexin V/7AAD assay using flow cytometer. Bar diagrams indicate the percentage of cell death with respective treated groups (Mean ±SEM, n=3). Two-way ANOVA. **G)** Indicated 5FU resistant OSCC lines and PDC2 cells were treated with 5 μM of 5FU and/or 50nM of Lestaurtinib for 48h, after which immunostaining was performed for γ-H2AX as described in materials ad methods. **H)** Indicated 5FU resistant OSCC lines and PDC2 cells were treated with Lestaurtinib for 48h, after which immunoblotting was performed with indicated antibodies. **I)** Effect on 5FU IC50 upon Lestaurtinib treatment in cells with or without MINK1 knockout in indicated 5FU resistant OSCC lines and PDC2 cells (n=3), \**P* < 0.05 by 2-way ANOVA.

**Figure 6:**
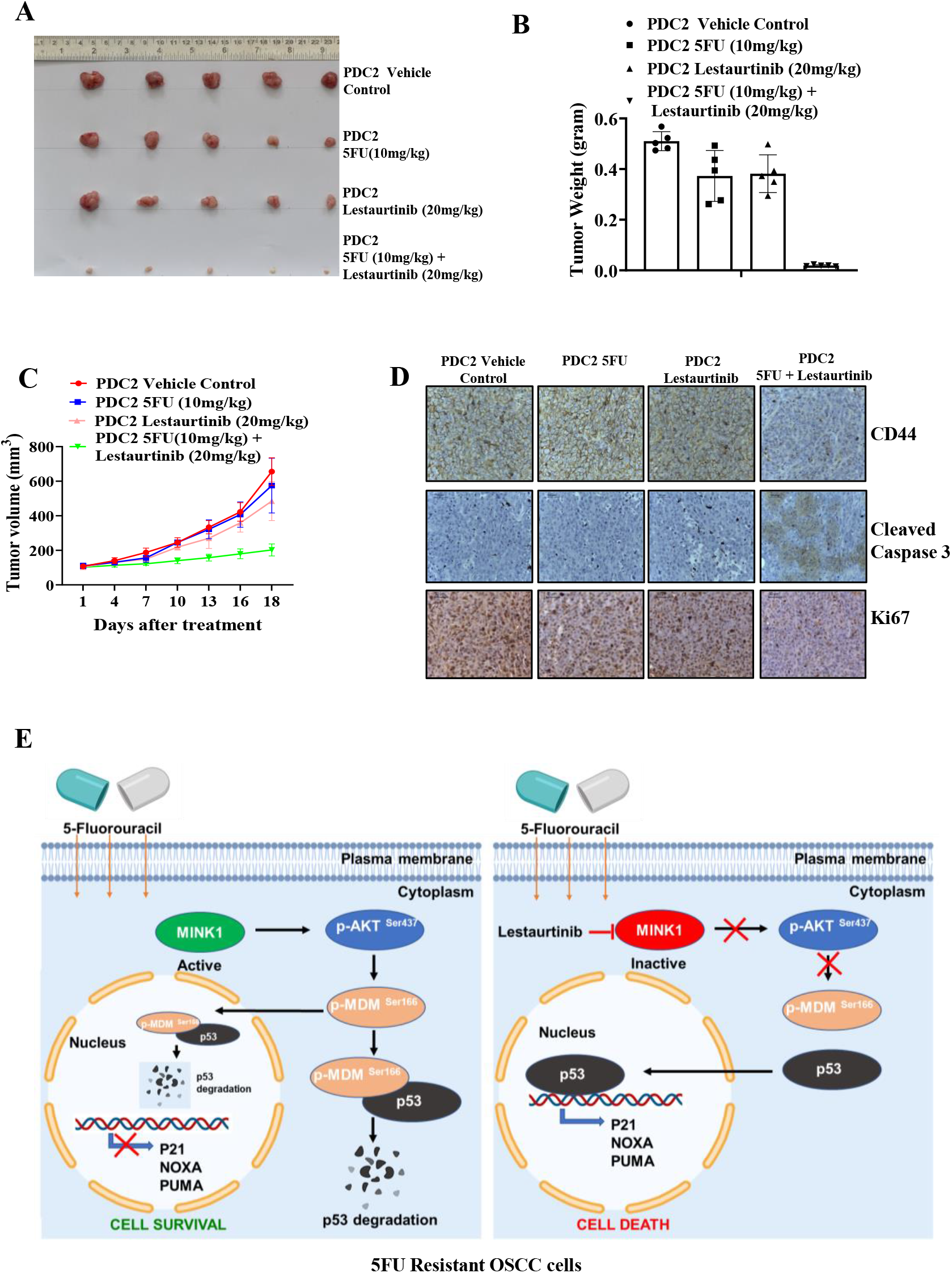
Lestaurtinib and 5FU synergistically reduced tumor burden *in vivo* in drug resistant OSCC: **A)** Patient-derived cells (PDC2) were earlier established from tumor of chemotherapy (TPF) non-responder patient. PDC2 were implanted in the right upper flank of athymic male nude mice, after which they were treated with 5FU and/or Lestaurtinib at indicated concentrations. At the end of the experiment mice were euthanized, and tumors were isolated and photographed (n = 5). **B)** Bar diagram indicates the tumor weight measured at the end of the experiment (mean ± SEM, n = 5). Two-way ANOVA. **C)** Tumor growth was measured at the indicated time points using digital slide caliper and plotted as a graph (mean ± SEM, n = 5). Two-way ANOVA. **D)** After completion of treatment, tumors were isolated, and paraffin-embedded sections were prepared as described in Methods to perform IHC with indicated antibodies. Scale bars: 50 μm. **E)** Schematic presentation of the mechanism by which MINK1 regulates p53 expression through AKT/MDM2 axis.

**Figure 7:**
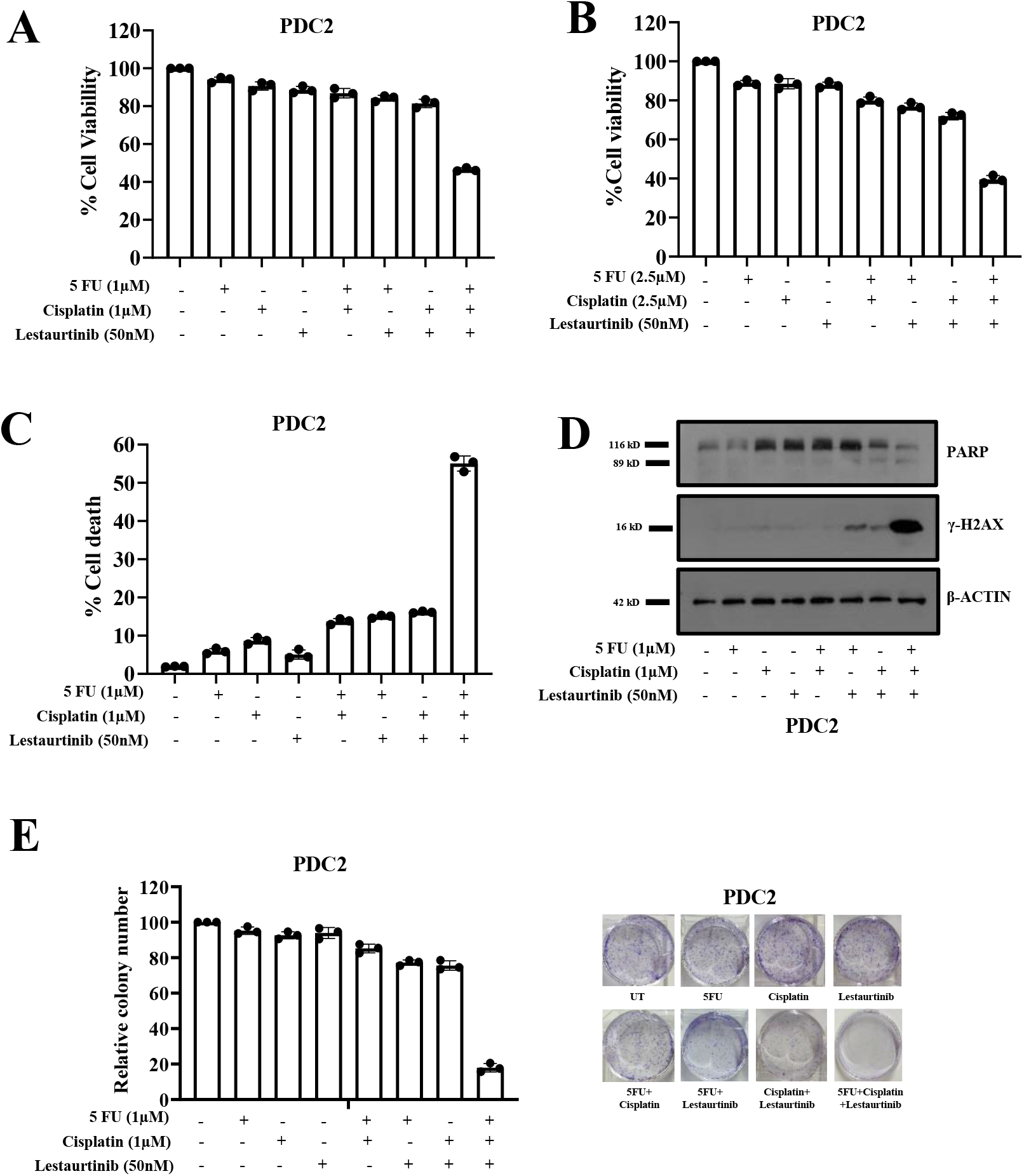
Evaluation of combinatorial anti-tumor effect of low dose of cisplatin, 5FU and lestaurtinb in TPF resistant patient derived cells (PDC2). **A-B)** PDC2 cells were treated with indicated concentrations of cisplatin, 5FU and lestaurtinib for 48h and cell viability was measured by MTT assay(n=3 and *P < 0.05 by 2-way ANOVA). **C)** PDC2 cells were treated with indicated concentrations of cisplatin, 5FU and lestaurtinib for 48h after which cell death was determined by annexin V/7AAD assay using flow cytometer. Bar diagrams indicate the percentage of cell death (early and late apoptotic) with respective treated groups (Mean ±SEM, n=3 by Two-way ANOVA). **D)** PDC2 cells were treated with indicated concentrations of cisplatin, 5FU and lestaurtinib for 48h and immunoblotting was performed with indicated antibodies. **E)** PDC2 cells were treated with indicated concentrations of cisplatin, 5FU, lestaurtinib for 12 days and colony forming assays were performed as described in method section. Left panel: Bar diagram indicate the relative colony number (n=3 and *P < 0.05 by 2-way ANOVA). Right panel: representative photographs of colony forming assay in each group.

## Discussion

The hallmark chemoresistant phenotypes of cancer cells are reduced apoptosis, altered metabolic activity, enhanced cancer stem cells like population, increased drug efflux and decreased drug accumulations. However, the causative factors which are responsible for acquired chemoresistance is largely not known. It is well known that kinases play key role in various processes of carcinogenesis and kinase inhibitors are established as potential anti-tumor agents. In this study, for the first time, we have performed a kinome screening in drug resistant cancer cells to explore the potential kinase(s) those mediate 5FU resistance in OSCC. From the primary and secondary kinome screening, MINK1 was found to be top ranked kinase that can re-sensitize drug resistant cells to 5FU. Overall, MINK1 is known to regulate cell senescence, cell motility and migration. Till date the potential role of MINK1 in modulating chemoresistanace is still unknown. Here in this study, we found the novel function of MINK1 by which it regulates 5FU resistance in OSCC.

To understand the mechanism by which MINK1 regulates 5FU resistance, we performed a high-throughput phosphorylation profiling in 5FUR cells stably expressing MINK1sgRNA. From this study, we found p-p53 (Ser33) and p-53(Ser15) to be significantly up-regulated in MINK1KO cells and p-AKT (Ser473) and p-MDM2 (Ser166) were found to be down-regulated in MINK1KO cells as compared to MINK1WT. The tumor suppressor p53 is phosphorylated at various amino acids by different kinases, which tightly regulates its stability ^23^. It is well known that MDM2 (a E3 ubiquitin ligase) acts as a negative regulator of p53. MDM2 forms a complex with p53 and facilitates the recruitment of ubiquitin molecules for its degradation ^24^. Earlier, it was established that insulin induced activated AKT (Ser473) phosphorylates MDM2 at Ser 166 and Ser 186, which can lead to MDM2 mediated proteasomal degradation of p53 in cytoplasm as well as in nucleus ^25, 26^. These events lead to blocking of p53-mediated transcription of genes those generally involve in apoptosis, cell cycle regulation and senescence. In addition to this, p53 is phosphorylated at Ser15 by ATM, DNA-PK and ATR in response to DNA damage ^27, 28, 29^. Hence, phosphorylation of Ser15 and Ser33 leads to activation and stabilization of p53 as they attenuate the MDM2 mediated degradation of p53 ^30, 31^. Overall, in this study we found that MINK1 regulates the expression of p53 through activation of AKT which in turns triggers p-MDM2 (Ser 166) (Figure 6 E).

Jin et al 2018 performed a kinome screening in cisplatin resistant cells to explore the potential kinases those confer cisplatin resistance in HNSCC. The data suggests that microtubule-associated serine/threonine-protein kinase 1 (MAST1) mediates cisplatin resistance in HNSCC by phosphorylating MEK1, triggering cRaf-independent activation of MEK1, which led to down regulation of BH3 only protein BIM. Jin et al 2018 also found that lestaurtinib to be a potent inhibitor of MAST1. Lestaurtinib successfully restores the cisplatin induced cell death in cisplatin resistant cells ^6^. Lestaurtinib not only inhibits MAST1 activity but also known as an inhibitor of JAK2, Trk and FLT3 ^32, 33^. In this study, we found that lestaurtinib inhibits activity of MINK1 and lestaurtinib can resensitize the drug resistant OSCC to 5FU. The most common chemotherapy regimen for OSCC is the combination of cisplatin, 5FU and Docetaxel (TPF). Finally, our data suggests that combination of extremely low dose of cisplatin (1μM), 5FU (1μM) and lestaurtinib (50nM) can overcome chemoresistance in OSCC (Fig. S8). Currently lestaurtinib alone or in combination with other chemotherapy drug is under clinical investigation (phase II) for patients having AML and it is well tolerated in human beings ^34^.

Overall, our data suggests that MINK1 is a mediator of 5FU resistance in OSCC. Besides this, though we have demonstrated that MINK1 negatively regulates P53 through AKT/MDM2 axis in 5FU resistant OSCC, the 5FU specificity of this MINK1 driven signalling cascade remains to be fully elucidated. Further, genetic or pharmacological (Lestaurtinib) inhibition of MINK1 successfully resensitized chemo resistant lines to 5FU. This novel combination of 5FU and Lestaurtinib needs further clinical investigation.

## Acknowledgements

PM is a CSIR-SRF, SM is UGC-SRF, SAA is a UGC-JRF.

## Author contribution

SM, PM, OS, MP, SAA and SP performed experiments, and analyzed the data, under the direction of R.D. RR, MS and S.K.M. performed part of experiments. SM, PM, RKS and R.D. designed experiments and supervised the study. R.D wrote the manuscript. All authors approved the final version.

