## SUPPLEMENTARY FIGURES for "CRISPR-based kinome-screening revealed MINK1 as a druggable player to rewire 5FU-resistance in OSCC through AKT/MDM2/p53 axis"

### Supplementary Figure 01

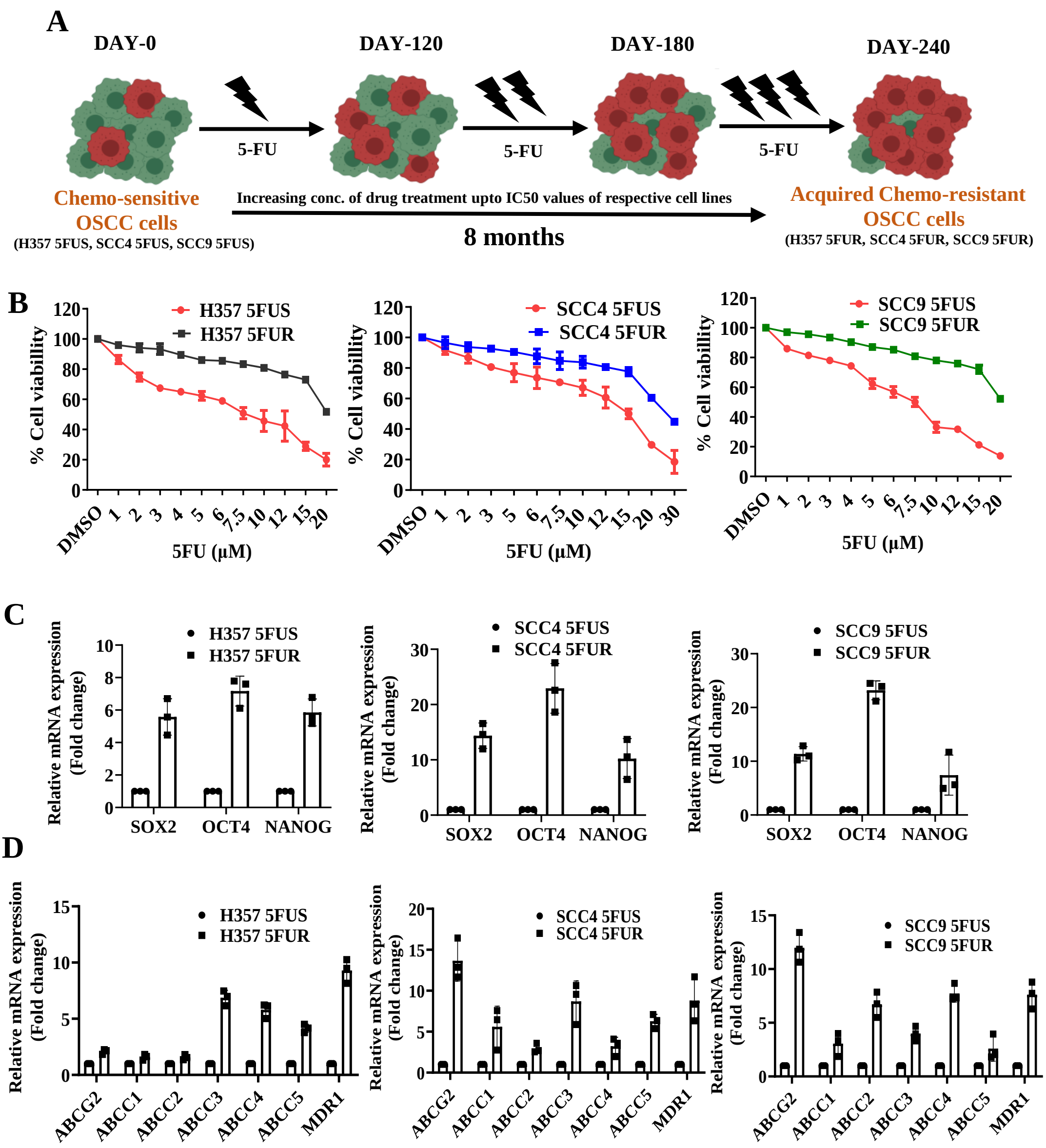

### Supplementary Figure 02

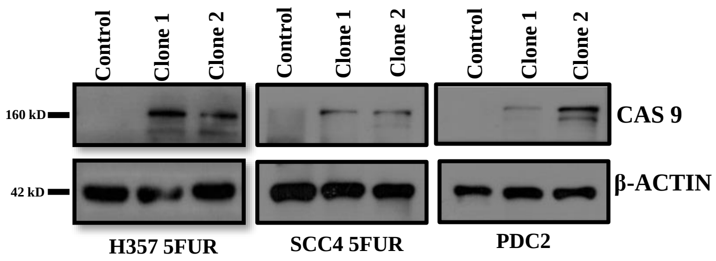

### Supplementary Figure 03

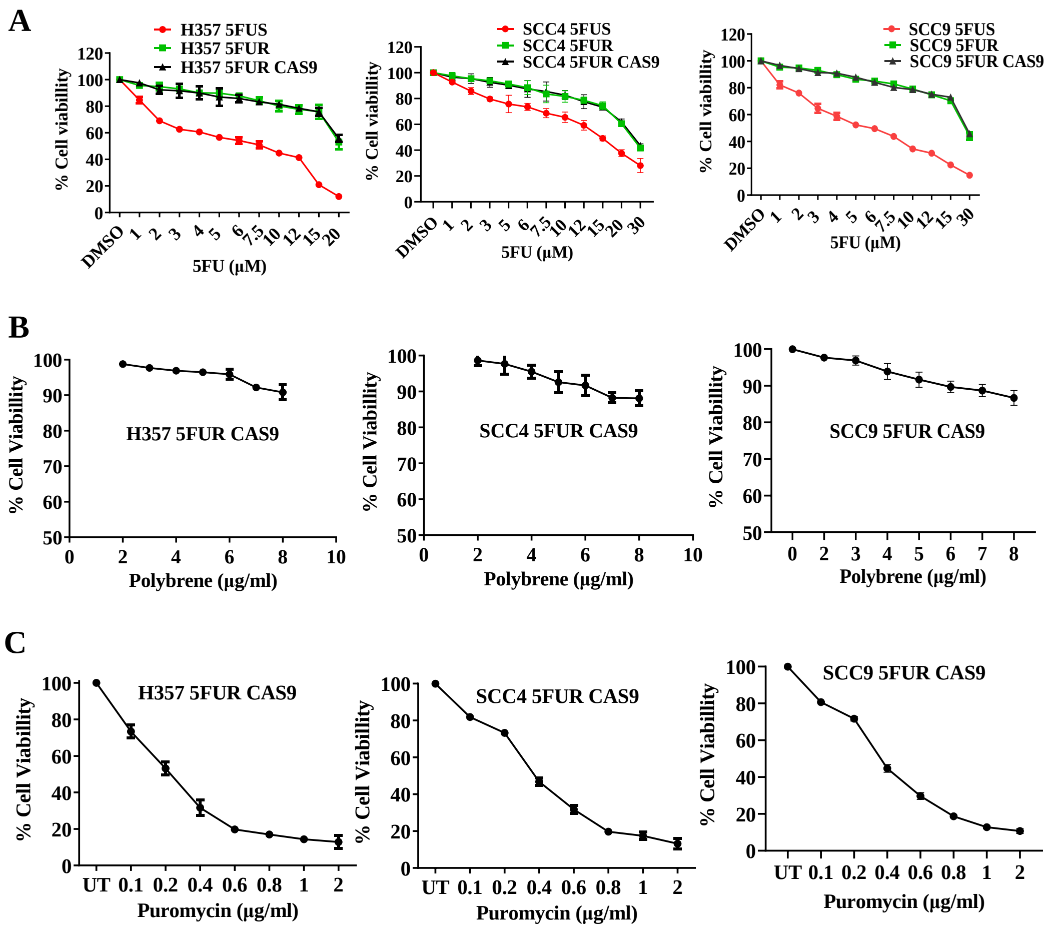

### Supplementary Figure 04

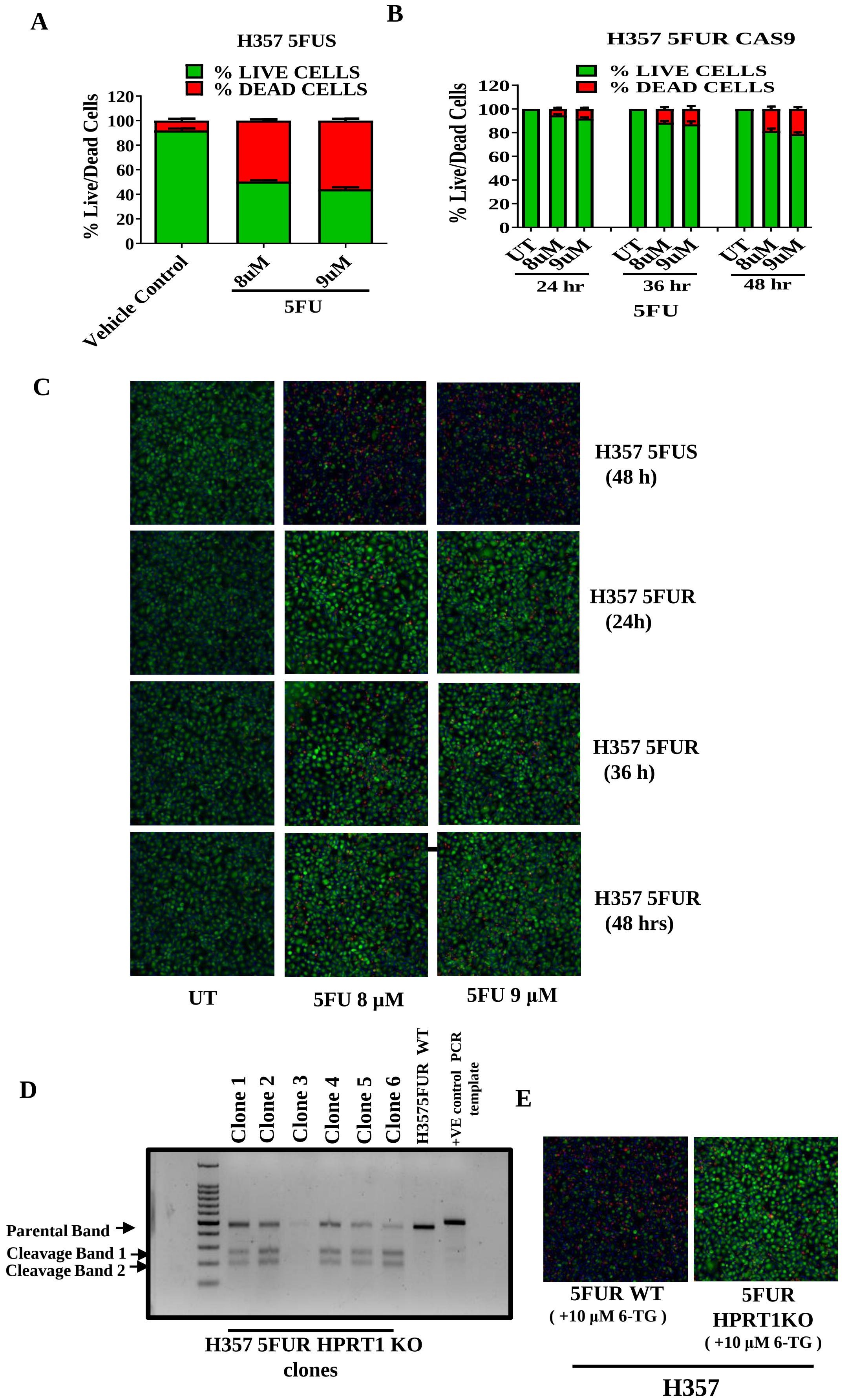

### Supplementary Figure 05

A

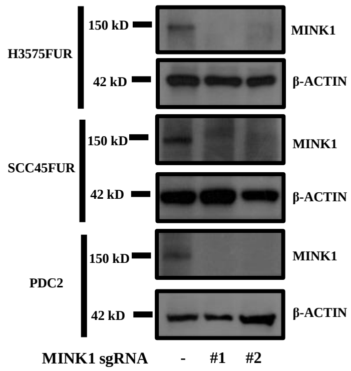


B

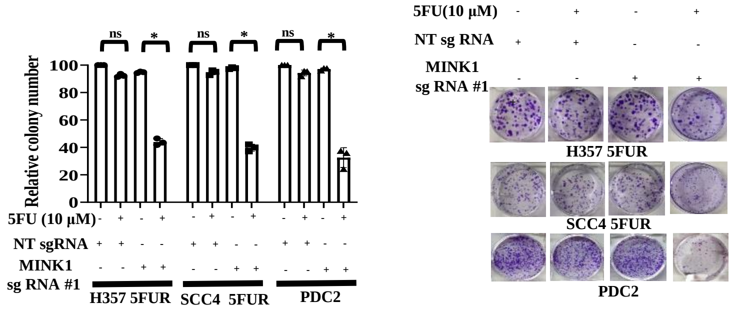


C

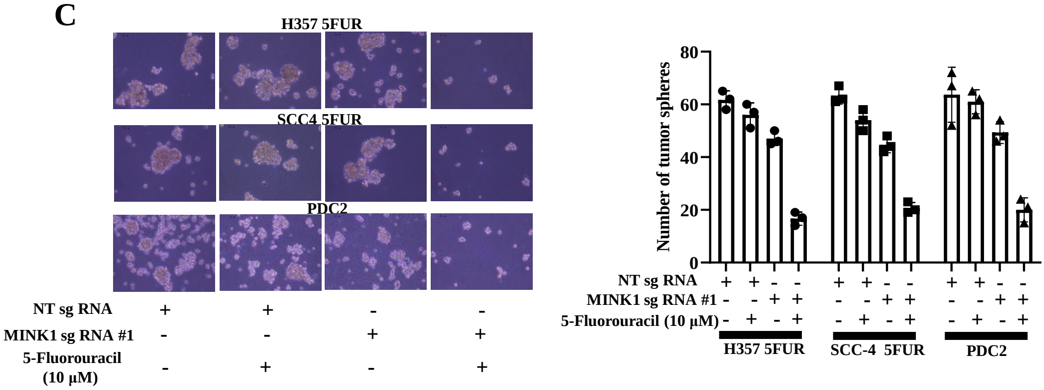

### Supplementary Figure 06

A

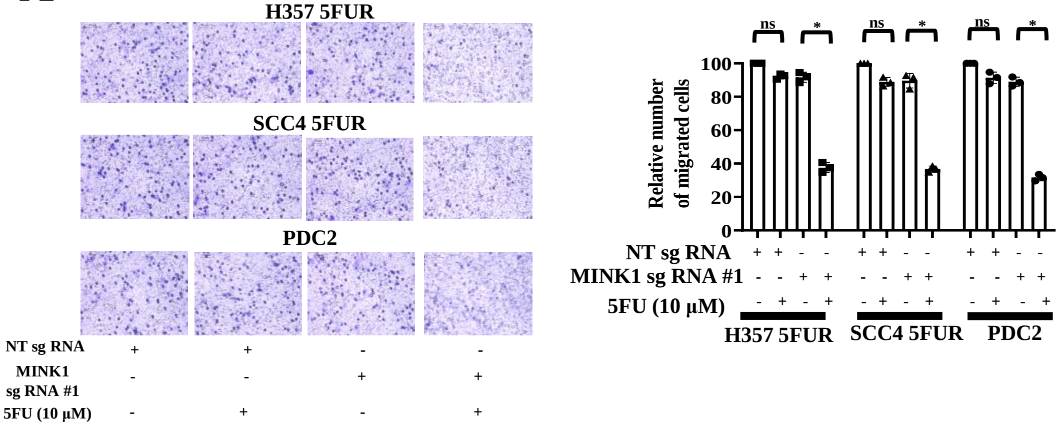


B

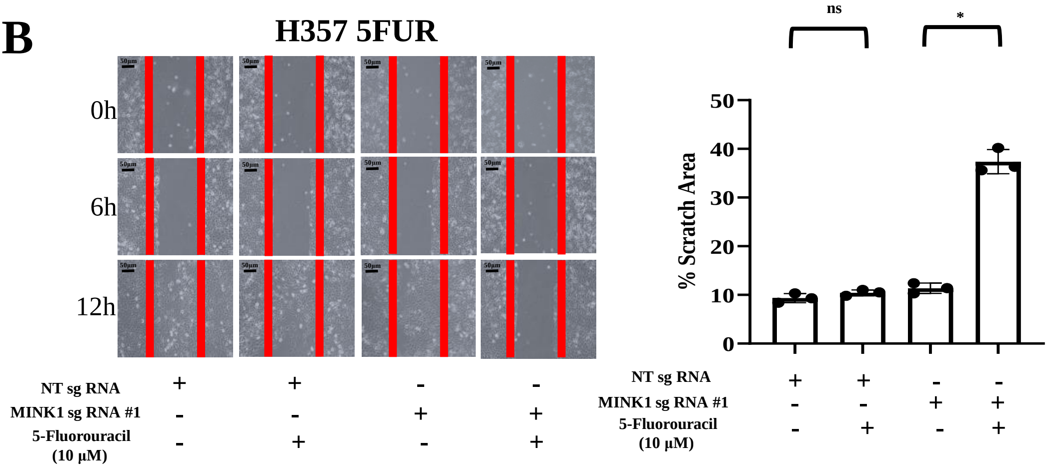


C

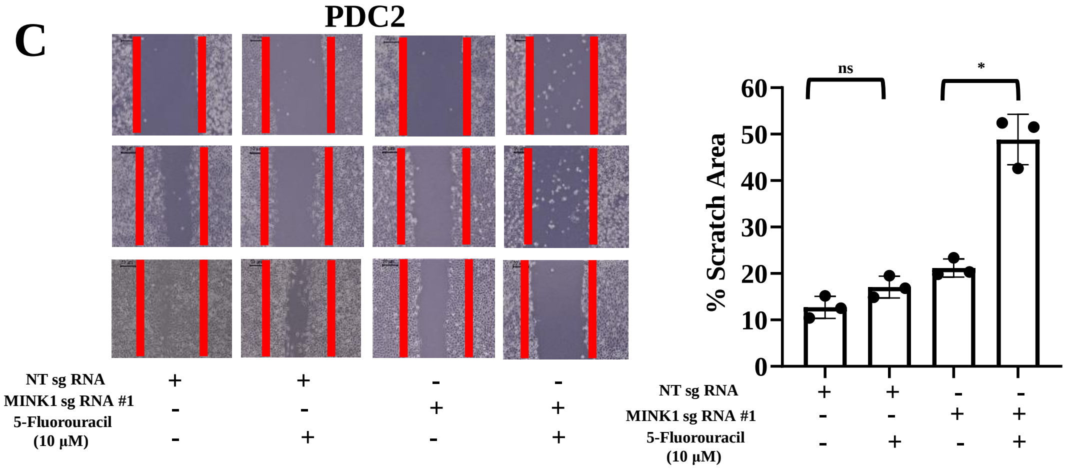

### Supplementary Figure 07

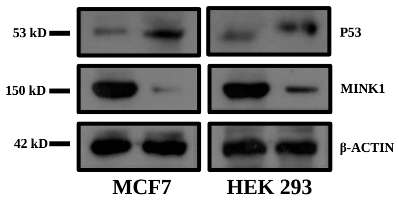

### Supplementary Figure 08

A

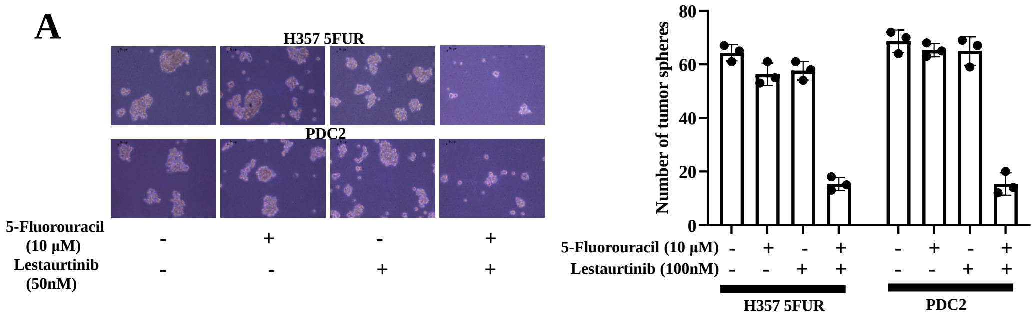


B

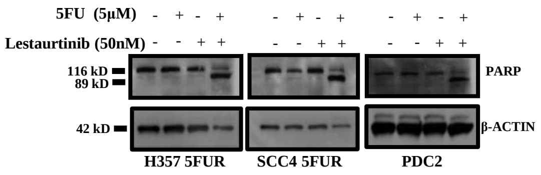


C

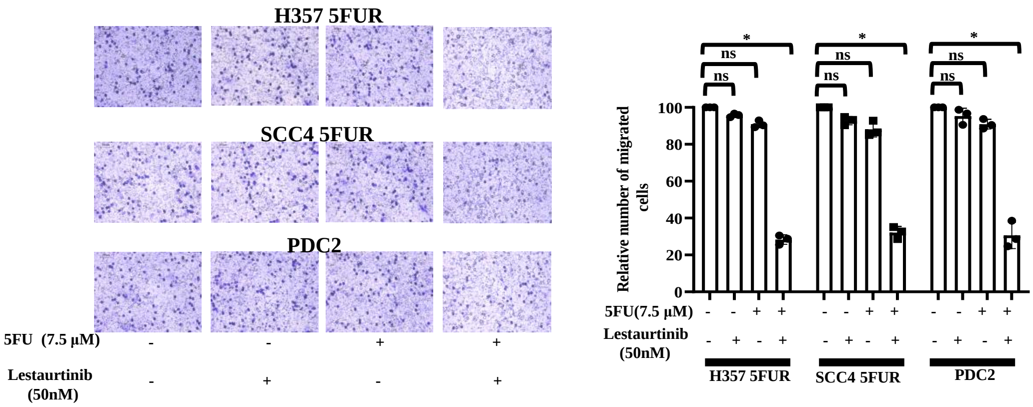
