## SUPPLEMENTARY FIGURES LEGEND for "CRISPR-based kinome-screening revealed MINK1 as a druggable player to rewire 5FU-resistance in OSCC through AKT/MDM2/p53 axis"

**Figure S1: Characterization of sensitive and 5FU resistant OSCC lines: A)** Schematic presentation of approach for establishing 5FU resistant OSCC lines **B)** Sensitive and 5FU resistant pattern (5FUS and 5FUR) of H357, SCC4 and SCC9 cells were treated with indicated concentrations of 5FU for 48h and cell viability was determined by MTT assay (n=3, *: $P < 0.05$). 2-way ANOVA. **C-D)** RNA was isolated from Sensitive and 5FU resistant pattern (5FUS and 5FUR) of H357, SCC4 and SCC9 cells and relative mRNA (fold change) expression of indicated genes was analyzed by qRT-PCR (mean ± SEM, n = 3), 2-way ANOVA.

**Figure S2: Overexpression of Cas9 in 5FU resistant OSCC lines and PDC: A)** Lysates were collected from indicated cells and immunoblotting was performed with indicated antibodies.

**Figure S3: Characterization of Cas9 overexpressing OSCC lines regarding 5FU resistance, polybrene tolerance and puromycin sensitivity: A)** Indicated sensitive, 5FU resistant, 5FU resistant Cas9 overexpressing cells were treated with indicated concentrations of 5FU for 48h and cell viability was determined by MTT assay (n=3 and *$P < 0.05$ by 2-way ANOVA). **B)** Indicated 5FU resistant Cas9 overexpressing cells were treated with indicated concentrations of polybrene for 48h and cell viability was determined by MTT assay (n=3 and *$P < 0.05$ by 2-way ANOVA). **C)** Indicated 5FU resistant Cas9 overexpressing cells were treated with indicated concentrations of puromycin for 48h and cell viability was determined by MTT assay (n=3 and *$P < 0.05$ by 2-way ANOVA).

**Figure S4: Validation of 5FU resistance and optimization of kinome screening conditions using high content analyzer: A-B)** 5FU sensitive and resistant patterns of H357 cells were treated with indicated concentrations of 5FU for the indicated time points, after which cell viability was measured in high content analyzer using a live/dead cell imaging kit. **C)** The representative fluorescent images acquired from high content analyzer with indicated treated groups in indicated cells. **D)** Different HPRT1 KO clones in H357 5FUR cells were subjected to genomic cleavage detection assay as described in materials and methods section. **E)** The representative fluorescent images acquired from high content analyzer with indicated treated groups in indicated cells.

**Figure S5: Targeting MINK1 reduced cell proliferation and stemness in chemoresistant OSCC: A)** MINK1 knock out clones were generated using a lentiviral approach expressing 2 different sgRNAs (#1 and #2) in Cas9 overexpressing 5FU resistant lines and PDC2. Immunoblotting was performed with indicated antibodies in indicated cells. Patient derived cells (PDC2) was established from tumor of TPF treated chemo-nonresponder patient. **B)** MINK1 KO and MINK1WT cells were treated with 5FU for 12 days and colony forming assays were performed as described in method section. Left panel: Bar diagram indicate the relative colony number (n=3 and *$P < 0.05$ by 2-way ANOVA). Right panel: representative photographs of colony forming assay in each group. **C)** Left panel: Tumor spheroid assay was performed as described in Methods with indicated cells stably expressing NTSgRNA and MINK1sgRNA#1 followed by treatment with indicated concentration of 5FU for 5 days. At the end of the experiment, spheroid photographs were captured using Leica DMIL microscope. Scale bar, 200 µm. Right panel: Number of tumor spheres formed from the experiment in A was counted. (n = 3), 2-way ANOVA.

**Figure S6: MINK1 genomic ablation negatively affects tumor cell migration in chemoresistant OSCC: A)** Indicated MINK1 WT and KO cells were treated with vehicle control or 5FU (10 µM) for 48h and were subjected to Boyden chamber assay as described in materials and methods section. Left panel: representative photographs of Boyden chamber assay in each group. Right panel: Bar diagrams indicate the relative number of migrated cells (n=3 and *$P$ < 0.05 by 2-way ANOVA). Scale bars: 50 µm. **B-C)** Indicated MINK1 WT and KO cells were treated with vehicle control or 5FU (10 µM) for 48h and were subjected to scratch assay as described in materials and methods section. Left panel : representative photographs of scratch assay in each group. Right panel : Bar diagrams indicate the percentage scratch area (n=3 and *$P$ < 0.05 by 2-way ANOVA). Scale bars: 50 µm.

**Figure S7: MINK1 regulates the expression of wild type p53:** Lysates were collected from indicated cells and immunoblotting was performed with indicated antibodies

**Figure S8: Lestaurtinib negatively affects stemness and tumor cell migration in chemoresistant OSCC: A)** Left panel: Tumor spheroid assay was performed as described in Methods with indicated cells with indicated concentration of 5FU and or lestaurtinb for 5 days. At the end of the experiment, spheroid photographs were captured using Leica DMIL microscope. Scale bar, 200 µm. Right panel: Number of tumor spheres formed from the experiment in A was counted. (n = 3), 2-way ANOVA.**B)** Indicated 5FU resistant OSCC lines and PDC2 cells were treated with 5FU and or Lestaurtinib for 48h, after which immunoblotting was performed with indicated antibodies. **C)** Indicated 5FU resistant OSCC lines and PDC cells were treated with vehicle control, 5FU (7.5 µM) and/or lestaurtinb (50 nM) for 48h, followed by subjecting to Boyden chamber assay. Left panel: representative photographs of Boyden chamber assay in each group. Right panel: Bar diagrams indicate the relative number of migrated cells (n=3 and *$P < 0.05$ by 2-way ANOVA). Scale bars: 50 μm.
