## SUPPLEMENTARY MATERIALS AND METHODS for "CRISPR-based kinome-screening revealed MINK1 as a druggable player to rewire 5FU-resistance in OSCC through AKT/MDM2/p53 axis"

**Generation of 5-Fluorouracil resistant cell lines:** For establishment of 5FU resistant cell lines, human OSCC cell lines (H357, SCC4 and SCC9) were initially treated with 1µM (lower dose) of 5-Fluorouracil for a week and then the concentration of 5FU was gradually increased up to the IC50 value, i.e. 10 µM for H357, 15 µM for SCC4 and 7.5 µM for SCC9 within a span of 3 months. Parental cells were grouped as sensitive (H357 5FUS, SCC4 5FUS and SCC9 5FUS) and after a period of 8 months of 5FU treatment, they were termed as 5FU Resistant (H357 5FUR, SCC4 5FUR and SCC9 5FUR) cells.

**Genomic Cleavage Detection Assay:** The genomic cleavage efficiency was measured using the GeneArt® Genomic Cleavage Detection kit (Thermo Fisher Scientific, Cat # A24372). For this, cells were lysed and DNA was extracted, followed by PCR amplification of the region on genome, where Cas9 endonuclease introduced a cleavage, using specifically designed primers as described in the protocol of the kit. Further, the PCR products were denatured and allowed for random reannealing, so that mismatches are generated as a result of genomic insertions or deletions (indels) created by the cellular repair mechanisms following the cleavage induced by Cas9. These mismatches were subsequently detected and cleaved by Detection Enzyme and then the resultant bands were analyzed by agarose gel electrophoresis. Primers used in this study for cleavage detection assay are mentioned in Table S2.

**Lentivirus production and generation of stable MINK1 KO cell lines** : LentiCas9-Blast was obtained from addgene (#52962) which is kindly deposited by Feng Zhang lab [12]. sgRNAs targeting MINK1 were cloned into pKLV2-U6gRNA5(BbsI)-PGKpuro2ABFP-W (Addgene #67974) vector as per the protocol mentioned in addgene, which is kindly deposited by Kosuke Yusa lab [13]. Respective lentiviruses were produced by transfection of LentiCas9-Blast or pKLV2-U6gRNA5(BbsI)-PGKpuro2ABFP-W plasmid along with packaging plasmid psPAX2 and envelop plasmid pMD2G into HEK293T cells as described in Shriwas et al [14]. Further 5FUR cells were infected with MINK1 sgRNA lentivirus using polybrene (8µg/ml) followed by puromycin (up to 5µg/ml) selection. After a week cells were picked up and seeded in 96 well plates with 1 cell/well dilution. Confirmations were done by western analysis. All sgRNA sequences used in this study are mentioned in Table S2.

**Lentivirus production and generation of stable MINK1 KD cell lines:** pLKO.1vector was obtained from addgene (Cat #10878), which is kindly deposited by David Root lab [15]. shRNAs targeting MINK1 were cloned into pLKO.1vector as per the protocol mentioned in addgene. Lentivirus was produced by transfection of pLKO.1 plasmid along with packaging plasmid psPAX2 and envelop plasmid pMD2G into HEK293T cells. Further 5FUR cells were infected with MINK1 shRNA lentivirus using polybrene (8µg/ml) followed by puromycin (up to 5µg/ml) selection. After 15 days colonies were picked up and confirmations were done by western analysis. All shRNA sequences used in this study are mentioned in Table S2.

**RT-PCR and Real Time Quantitative PCR**: RNA mini kit (Himedia, Cat# MB602) was used to isolate total RNA as per manufacturer's instruction and quantified by Nanodrop. Verso cDNA synthesis kit (ThermoFisher Scientific, Cat # AB1453A) was used to synthesize c-DNA by reverse transcription PCR using 300 ng of RNA. qRT-PCR was carried out using SYBR Green master mix (Thermo Fisher scientific Cat # 4367659). GAPDH was used as a loading control. The primers (oligos) sequence used for qRT-PCR in this study are listed in Table S2.

**Immunoblotting:** Immunoblotting was performed by loading equal amounts of cell lysates as described earlier [16]. In this study, primary antibodies used were against β-actin (Sigma, Cat#A2066), Cas9 (CST, Cat #14697S), MINK1 (Sigma, Cat# HPA056296, Invitrogen, Cat# PA5-28901), P53 (CST, Cat #2527T), p-P53(Ser15) (CST, Cat #9284T), p-P53(Ser33) (CST,

Cat #2526), PARP (CST, Cat #9542L), $p^{s-139}$-H2AX (CST, Cat # 9718S), AKT (CST, Cat #9272S), pAKT(Ser473) (CST, Cat #4058S), MDM2 (Santa Cruz, Cat #sc965), pMDM2 (ser166) (Abcam, Cat # ab131355), TIGAR (Abcam, Cat # Ab37910), P21(CST, Cat # 2947S), NOXA (Imgenex, Cat # IMG-349A).

**Assessment of cell viability:** Cell viability was measured by 3-(4, 5-dimethylthiazol-2-yl)-2, 5-diphenyltetrazolium bromide (MTT; Sigma-Aldrich) assay as per manufacturer's instruction.

**Colony formation assay**: Colony formation assay was performed as described in Shriwas et al [17].

**Annexin-V PE/7-AAD Assay**: Apoptosis and cell death assay was performed by using Annexin V Apoptosis Detection Kit PE (eBioscience™, USA, Cat # 88-8102-74) as described earlier [17] and cell death was monitored using a flow cytometer (BD FACS Fortessa, USA).

**Tumorsphere formation assay:** Tumorsphere assay was performed as described in Mohapatra et al [18]

**Immunofluorescence:** The cells were seeded on lysine coated coverslip and cultured for overnight. On next day, cells were treated with 5FU for 48 hrs followed by 4% formaldehyde fixation for 15 mins. Next, cells were permeabilized with 1 × permeabilization buffer (eBioscience 00-8333-56) for 45 mins, followed by blocking with 3% BSA for 1 h at room temperature. After which, the cells were incubated with primary antibody overnight at 4 °C, washed three times with PBST pH 8.0 followed by 1hr incubation with Goat anti–Rabbit IgG(H+L) secondary Antibody, Alexa Fluor® 488 conjugate (Invitrogen, Cat #A -11008). After washing three times with PBST pH 8.0, cells were mounted with DAPI (Slow Fade ® GOLD Antifade, Thermo Fisher Scientific, Cat # S36938). Images were captured using a confocal microscopy (LEICA TCS-SP8). Anti $p^{s-139}$-H2AX (CST, Cat # 9718S) primary antibody was used for this study.

**Immunohistochemistry:** Immunohistochemistry of formalin fixed paraffin-embedded samples (OSCC patients' tumors and Xenograft tumors from mice) were performed as described previously [18]. Antibodies against MINK1 (Sigma, Cat# HPA056296, Invitrogen, Cat# PA5-28901), CD44 (NOVUS, Cat# NBP1-31488), Cleaved Caspase-3 (CST, Cat# 9661S), Ki67 (Vector, Cat #VPRM04) were used for IHC. Images were obtained using Leica DM500 microscope. Q-score was calculated by multiplying percentage of positive cells with staining (P) and intensity of staining (I). P was determined by the percentage of positively stained cells in the section and I was determined by the intensity of the staining in the section i.e. strong (value=3), intermediate (value=2), weak (value=1) and negative (value=0).

**Transient transfection and overexpression of MINK1 and myr-AKT in MINK1 KD cell lines:** pDONR223-MINK1 (Plasmid #23522) was procured from addgene followed by transfer of the insert into pLenti CMV/TO Puro DEST (670-1) (Addgene, Cat#17293) destination vector by gateway cloning method using Gateway LR Clonase II Plus Enzyme mix (Invitrogen, Cat# 1756069). These plasmids were kindly deposited by William Hahn, David Root labs [19] and Eric Campeau, Paul Kaufman labs [20] in addgene. Further, MINK1 knockdown cells, stably expressing shRNA#1 targeting 3' UTR of MINK1 mRNA, were transiently transfected with pLenti CMV/TO Puro DEST-MINK1 using the ViaFect transfection reagent (Promega Cat# E4982). The transfection efficiency was confirmed by immunoblotting against Anti-MINK1. pLNCX myr HA Akt1 (Addgene, Cat #9005) was used for transient overexpression of constitutively activated AKT. The myr HA Akt1 vector was kindly deposited to Addgene by Sellers WR lab [21].

**OSCC patient sample:** Loco regionally advanced OSCC samples were collected. Neoadjuvant chemotherapy has been prescribed before surgery and/or radiotherapy. The three-drug combination of TPF is having highest response (TAX 324). After chemotherapy (CT) the response is evaluated as per RECIST criteria (Response evaluation criteria in solid tumors) by clinical and radiological evaluation. After chemotherapy the patient can be grouped as complete response (CR), Partial Response (PR), stable disease (SD) or Progressive Disease (PD). If there was no evidence of malignancy, then it was diagnosed as complete response (CR). If the target lesions had decreased more than equal to 30% of the sum of the longest diameter, then it was diagnosed as partial response (PR). If there was no sign of either CR or PR, then it was called stable disease, and if the target lesions had increased more than or equal to 20% of the sum of the longest diameter, then it was called PD (progressive disease). As the patients showing CR and PR have responded to the CT they are categorized as Responders and the patients with stable disease or PD with almost no response to CT are categorized as Non-Responders. Human Ethics Committee (HEC) of the Institution of Life Sciences approved all patient-related studies, and informed consent was obtained from all patients. Study subject details with treatment modalities are presented in Table S1a and b.

**Zebrafish xenograft:** The experimental protocols used for this work were approved by the institutional animal ethical review committee (ILS/IAEC-214-AH/APR-21). PDC2 control and MINK1 knockdown stable cells were suspended in individual tubes at a density of 1x $10^6$ cells/ml in normal media followed by addition of 5μl of the cell-labeling solution (Vybrant™ DiI Cell-Labeling Solution Catalog number: V22885) and mixed well by gentle pipetting. The cells were incubated for 20 minutes at 37°C to obtain uniform labeling and then centrifuged at 1100 rpm for 5 minutes at RT. The supernatant was removed and cells were resuspended gently in 1ml of warm (37°C) media and centrifuged at 1100 rpm for 2 minutes for the removal of extra dye and the wash step was repeated twice. The Dil stained cells were resuspended at final density 200 cells/nl and ~400 cells were microinjected (Femtojet microinjector) into perivitelline space of 48 hpf embryos of zebrafish (*Danio rerio*) [Tg(fli1:EGFP)] for the development of tumor. The images of zebrafish embryos were captured using a fluorescence stereomicroscope (Leica MZ16) on the day of injection (Day 0), followed by 5FU treatment

(500μM) 3 days post injection (Day 3) and then final imaging 5 days after injection (Day 5). The tumor growth was assessed by an increase or decrease in fluorescence intensity on the 5th day compared to the day of injection. The quantitation of fluorescence intensity was performed using ImageJ software and represented as mean fluorescent intensity where day 0 readings was taken as baseline.

**Trans-well migration assay**: For trans-well migration assay, MINK1WT and MINK1 KO chemoresistant cells were treated with vehicle control or 5FU (10 μM, 48hrs). In another experimental set up, 5FU resistant cells were treated with vehicle control, 5FU (10 μM), Lestaurtinib (50 nM), or 5FU (10 μM) and Lestaurtinib (50 nM). After treatment, cells were seeded at a density of $1 \times 10^4$ cells in the upper chamber of 24-well trans-well system in 250 μL of serum free DMEM-F12 medium. 750 μL of DMEM-F12 Medium supplemented with 10% serum, used as a chemo-attractant, was added in the lower chamber. After 24 hrs of incubation, the cells inside the upper chamber were scrubbed with a cotton swab, whereas the migrated cells were fixed and stained with crystal violet, followed by quantification of migration by manual counting using a microscope.

***In vitro* scratch assay:** *In vitro* scratch assay was performed as described earlier [16].
