## SUPPLEMENTARY TABLES for "CRISPR-based kinome-screening revealed MINK1 as a druggable player to rewire 5FU-resistance in OSCC through AKT/MDM2/p53 axis"

### Table S1a

#### chemotherapy-responder patient details

| Sl No | Tumor samples | Age/Sex | Site of disease | Clinical stage | Chemotherapy (NACT) | Cycle |
|---|---|---|---|---|---|---|
| 1 | **Patient#1** | 42/M | Tongue Rt lateral border | T4aN1M0 | Docetaxel + Cisplatin+ 5FU | 2 |
| 2 | **Patient#2** | 67/M | Tongue Lt lateral border | T4aN1Mx | Docetaxel + Cisplatin+ 5FU | 3 |
| 3 | **Patient#3** | 50/M | Rt- Buccal mucosa | T4aN2bM0 | Docetaxel + Cisplatin+ 5FU | 3 |
| 4 | **Patient#4** | 75/M | Oral cavity | T3N2bM0 | Doceaqualip + Carboplatin | 3 |
| 5 | **Patient#5** | 46/M | Tongue | T3N1M0 | Docetaxel + Cisplatin+ 5FU | 3 |
| 6 | **Patient#6** | 35/M | Tongue | T4aN2eM0 | Docetaxel + Cisplatin+ 5FU | 3 |
| 7 | **Patient#7** | 38/M | Right Buccal Mucosa | T4bN2bM0 | Docetaxel + Cisplatin+ 5FU | 3 |
| 8 | **Patient#8** | 34/M | Left Buccal Mucosa | T4aN2bMx | Docetaxel + Cisplatin+ 5FU | 3 |
| 9 | **Patient#9** | 40/M | Tongue | T2N2cM0 | Docetaxel + Cisplatin+ 5FU | 3 |
| 10 | **Patient#10** | 45/M | Tongue | T2N1M0 | Docetaxel + Cisplatin+ 5FU | 3 |
| 11 | **Patient#11** | 51/M | Tongue Lt. lateral border | T4aN2cM0 | Docetaxel + Cisplatin+ 5FU | 3 |

Chemotherapy Doses: **Cisplatin**: 100mg. **Docetaxel**: 100mg, **5FU**:1000mg, **Doceaqualip**: 80mg, **Carboplatin:** AUC 4 (area under the ROC curve)

### Table S1b

#### Chemotherapy-non-responders patient Details

| Sl No | Tumor samples | Age /Sex | Site of disease | Clinical stage | Chemotherapy (NACT) | Cycle |
|---|---|---|---|---|---|---|
| 1 | **Patient# 1** | 76/M | Tongue Rt lateral border | T4N0M0 | Paclitaxel + Cisplatin | 3 |
| 2 | **Patient#2 (PDC#2)** | 51/M | Rt- Buccal mucosa | T2N2bM0 | Docetaxel + Cisplatin+ 5FU | 2 |
| 3 | **Patient# 3** | 60/M | Tongue Rt lateral border | T3N1M0 | Paclitaxel + Cisplatin +5FU | 3 |
| 4 | **Patient#4** | 33/M | Rt- Lower Alveolar mucosa | T3N1Mx | Docetaxel + Cisplatin | 3 |
| 5 | **Patient#5** | 60/F | Tongue Lt lateral border | T4N0M0 | Docetaxel + Cisplatin+ 5FU | 3 |
| 6 | **Patient#6** | 59/M | Tongue Rt lateral border | T4aN1M0 | Docetaxel + Cisplatin+ 5FU | 3 |
| 7 | **Patient#7** | 46/M | Tongue | T4N3M0 | Docetaxel + Cisplatin+ 5FU | 3 |
| 8 | **Patient#8** | 55/F | Rt- Buccal Mucosa | T4aN2M0 | Docetaxel + Cisplatin+ 5FU | 2 |
| 9 | **Patient#9** | 37/M | Tongue | T4N3M0 | Docetaxel + Cisplatin+ 5FU | 2 |
| 10 | **Patient#10** | 27/M | Lt-Buccal Mucosa | T4N2M0 | Docetaxel + Cisplatin+ 5FU | 2 |
| 11 | **Patient#11** | 46/F | Rt- oral cavity | T4N1M0 | Docetaxel + Cisplatin+ 5FU | 3 |
| 12 | **Patient#12** | 42/M | Rt- Buccal Mucosa | TxN3bM0 | Paclitaxel + Cisplatin+ 5FU | 2 |
| 13 | **Patient#13** | 30/M | Tongue Rt lateral border | T2N0Mx | Paclitaxel + Cisplatin+ 5FU | 3 |
| 14 | **Patient#14** | 52/M | Rt- Buccal Mucosa | T4N2M0 | Docetaxel + Cisplatin+ 5FU | 3 |
| 15 | **Patient#15** | 32/M | Tongue | T3N1M0 | Docetaxel + Cisplatin+ 5FU | 3 |
| 16 | **Patient#16** | 35/M | Tongue | T4aN2aM0 | Docetaxel + Cisplatin+ 5FU | 3 |
| 17 | **Patient#17** | 36/M | Left Buccal Mucosa | T4aN2aM0 | Docetaxel + Cisplatin+ 5FU | 3 |
| 18 | **Patient #18** | 38/M | | T4bN2bMO | Docetaxel + Cisplatin+ 5FU | 2 |
| 19 | **Patient # 19** | 55/M | Tongue, Left lateral border, | | Docetaxel + Cisplatin+ 5FU | 3 |
| 20 | **Patient # 20** | 35/M | Tongue | T4aN2eM0+ | Docetaxel + Cisplatin+ 5FU | 3 |
| 21 | **Patient # 21** | 36/M | Left Buccal Mucosa | cT4aN2aM0 | Docetaxel + Cisplatin+ 5FU | 3 |
| 22 | **Patient # 22** | 39/M | Right mandible | cT4bN0Mx | Docetaxel + Cisplatin+ 5FU | 3 |
| 23 | **Patient # 23** | 55/M | Tongue, Left lateral border, | cT4aN2cM0 | Docetaxel + Cisplatin+ 5FU | 3 |

Chemotherapy taken, but after 1-2 cycles became non responded.
**Chemotherapy Doses**: **Cisplatin**: 100mg. **Paclitaxe**l: 260 mg, **Docetaxel**: 100mg, **5FU**:1000mg **Lt-Left, Rt-Right**

PDC2: patient derived cells isolated from indicated OSCC patients.

#### Table S2

##### Oligos for SgRNA and ShRNA

| Sg and Sh RNA primers | Oligo sequence |
|---|---|
| MINK1 sg RNA F (SgRNA#1) | CACCCGCAACATCGCCACCTACTA |
| MINK1 sg RNA R (SgRNA#1) | AAACTAGTAGGTGGCGATGTTGCG |
| MINK1 sg RNA F (SgRNA#2) | CACCGTGGTCGGCAATGGAACCTA |
| MINK1 sg RNA R (SgRNA#2) | AAACTAGGTTCCATTGCCGACCAC |
| MINK1 3' UTR sh RNA F (ShRNA#1) | CCGGAATGTAGTGGCCTTGGATATCCTCGAGGATATCCAAGGCCACTACATTTTTTG |
| MINK1 3' UTR sh RNA R (ShRNA#1) | AATTCAAAAAAATGTAGTGGCCTTGGATATCCTCGAGGATATCCAAGGCCACTACATT |

##### Oligos for qRT-PCR

| qRT PCR Primers | Primer sequence |
|---|---|
| 18S qRT F | GTAACCCGTTGAACCCCATT |
| 18S qRT R | CCATCCAATCGGTAGTAGCG |
| GAPDH qRT F | TCGGAGTCAACGGATTTGGT |
| GAPDH qRT R | TTGCCATGGGTGGAATCATA |
| OCT4 qRT F | CGACCATCTGCCGCTTTGAG |
| OCT4 qRT R | CCCCCTGTCCCCCATTCCTA |
| SOX2 qRT F | CACCTACAGCATGTCCTACTC |
| SOX2 qRT R | CATGCTGTTTCTTACTCTCCTC |
| Nanog qRT F | CAACTGGCCGAAGAATAGCA |
| Nanog qRT R | GCAGGAGAATTTGGCTGGAA |
| ABCC1 qRT F | AGTGGAACCCCTCTCTGTTTAAG |
| ABCC1 qRT R | CCTGATACGTCTTGGTCTTCATC |
| ABCC2 qRT F | AATCAGAGTCAAAGCCAAGATGCC |
| ABCC2 qRT R | TAGCTTCAGTAGGAATGATTTCAGGAGCAC |
| ABCC3 qRT F | TCCTTTGCCAACTTTCTCTGCAACTAT |
| ABCC3 qRT R | CTGGATCATTGTCTGTCAGATCCGT |
| ABCC4 qRT F | TGATGAGCCGTATGTTTTGC |
| ABCC4 qRT R | CTTCGGAACGGACTTGACAT |
| ABCC5 qRT F | AGAGGTGACCTTTGAGAACGCA |
| ABCC5 qRT R | CTCCAGATAACTCCACCAGACGG |
| ABCG2 qRT F | CCGCGACAGTTTCCAATGACCT |
| ABCG2 qRT R | GCCGAAGAGCTGCTGAGAACTGTA |
| MDR1 qRT F | AGGAAGCCAATGCCTATGACTTTA |
| MDR1 qRT R | CAACTGGGCCCCTCTCTCTC |

##### Oligos to detect genomic cleavage detection assays

| | |
|---|---|
| HPRT1 GCD F | TACACGTGTGAACCAACCCG |
| HPRT1 GCD R | GTAAGGCCCTCCTCTTTTATTT |
